# Computational modelling of chromosome re-replication in mutant strains of fission yeast

**DOI:** 10.1101/2020.09.24.311449

**Authors:** Béla Novák, John J Tyson

## Abstract

Typically cells replicate their genome only once per division cycle, but under some circumstances, both natural and unnatural, cells synthesize an overabundance of DNA, either in a disorganized fashion (‘over-replication’) or by a systematic doubling of chromosome number (‘endoreplication’). These variations on the theme of DNA replication and division have been studied in strains of fission yeast, *Schizosaccharomyces pombe*, carrying mutations that interfere with the function of mitotic cyclin-dependent kinase (Cdk1:Cdc13) without impeding the roles of DNA-replication licensing factor (Cdc18) and S-phase cyclin-dependent kinase (Cdk1:Cig2). Some of these mutations support endoreplication, and some over-replication. In this paper, we propose a dynamical model of the interactions among the proteins governing DNA replication and cell division in fission yeast. By computational simulations of the mathematical model, we account for the observed phenotypes of these re-replicating mutants, and by theoretical analysis of the dynamical system, we provide insight into the molecular distinctions between over-replicating and endoreplicating cells. In case of induced over-production of regulatory proteins, our model predicts that cells first switch from normal mitotic cell cycles to growth-controlled endoreplication, and ultimately to disorganized over-replication, parallel to the slow increase of protein to very high levels.

## Introduction

The eukaryotic cell cycle is characterized by strict alternation of DNA replication during S phase and chromosome segregation in mitosis (M phase). During mitotic cell divisions, each S phase is followed by mitosis and every M phase is preceded by the replication of all chromosomes. Both of these chromosomal events are triggered by periodic activation of Cyclin-dependent kinases (Cdk) in complex with their regulatory subunits, cyclins. Most eukaryotic cells use different Cdk:cyclin pairs to trigger DNA replication and entry into mitosis (Morgan, 2007). The lower eukaryote, *Schizosaccharomyces pombe* (‘fission yeast’) uses the same Cdk (called Cdk1) in both complexes; Cdk1 associates with S phase cyclin (Cig2) to initiate DNA replication, while its complex with mitotic cyclin (Cdc13) brings about M phase (Nurse, 1990; Mondesert *et al*., 1996). The activities of Cdk1:Cig2 and Cdk1:Cdc13 oscillate during the cell cycle, with peaks during DNA replication and mitosis, respectively (Figure 1A). The alternating oscillations of Cdk1:Cig2 and Cdk1:Cdc13 generate a steadily increasing activity of Cdk1 (the sum of Cig2-and Cdc13-dependent kinase activities) from the beginning of DNA replication until the end of M phase (Fisher and Nurse, 1996; Stern and Nurse, 1996). At the end of mitosis Cdk1:Cdc13 activity is reduced to a value close to zero by proteasomal degradation of Cdc13, triggered by abrupt polyubiqutinylation by an E3 ligase, the Anaphase Promoting Complex/Cyclosome. Subsequently, a new batch of Cig2 is synthesized, and Cdk1:Cig2 activity peaks during the next S phase. Therefore, the strict alternation between S and M phases of the mitotic cycle in fission yeast is strongly coupled to the alternating peaks of Cdk1:Cig2 and Cdk1:Cdc13 oscillations.

**Figure 1:**
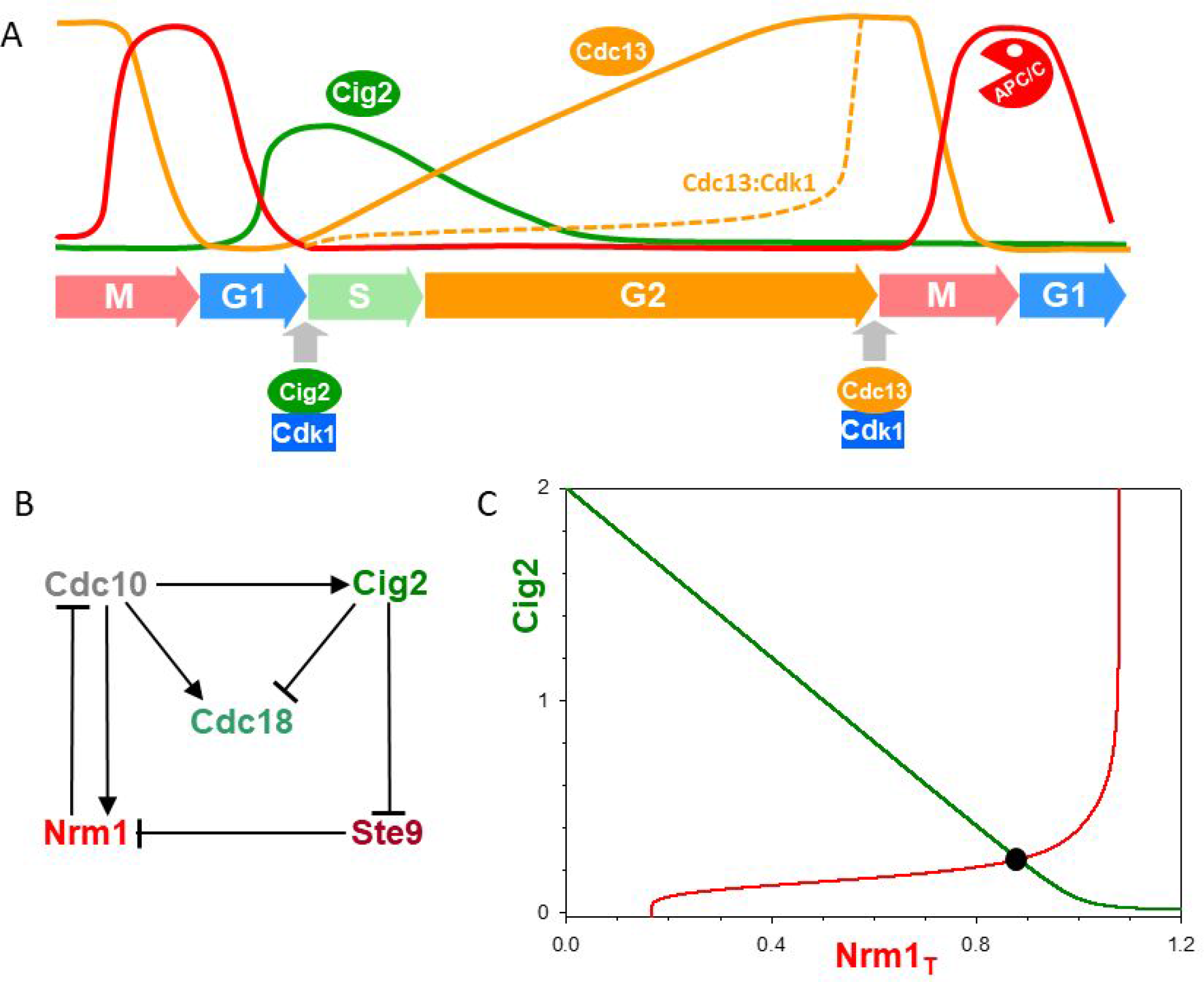
The core transcriptional negative feedback of fission yeast. (A) Fluctuation of cell cycle regulators during wild-type fission yeast cell cycle (schematic). (B) Influence diagram for Cdc10 downstream targets. (C) The negative feedback on the Nrm1_T_-Cig2 plane. Each dynamic variable is in steady state along its balance curve. Since Nrm1 inhibits the transcription of Cig2, Cig2 level is decreasing with Nrm1_T_ levels. Cig2 inhibits Nrm1 degradation, therefore Nrm1_T_ level shows a switch-like increase with Cig2. At the intersection of the two balance curves, the whole network is in steady state.

If an essential chromosomal event (e.g., DNA replication or mitotic spindle formation) becomes compromised, then further progression through the cell cycle must be suspended until the problem can be resolved. These potential problems are monitored by surveillance mechanisms that, when triggered, block the activation of the kinase complex (Cdk1:Cig2 or Cdk1:Cdc13) responsible to initiate the next event in the chromosome replication/division cycle. In addition, the oscillations of both Cdk1:cyclin complexes are controlled by a ‘cell size’ checkpoint. The critical size threshold permissive for oscillation is higher for the M-phase kinase than for the S-phase kinase (Wood and Nurse, 2015).

Interestingly, the normally interlinked oscillations of Cig2-and Cdc13-dependent kinases can run independently from each other, but these independent Cdk1 oscillations have different physiological outcomes. Since Cdc13 is the only essential cyclin for the fission yeast cell cycle, it can, in the absence of S phase cyclin (i.e., in *cig2*Δ cells), drive both DNA replication and mitosis (Fisher and Nurse, 1996; Coudreuse and Nurse, 2010). Since the activity of Cdk1 required for S phase is lower than for M phase (Coudreuse and Nurse, 2010), the oscillating activity of Cdk1:Cdc13 guarantees that initiation of DNA replication always precedes mitosis. Because the rate of synthesis of Cdc13 is uniform throughout the cell cycle, it is an underlying proteolytic negative feedback loop (through APC/C) that is responsible for the oscillation of Cdc13 level. The amplitude of the Cdk1:Cdc13 activity oscillation is boosted by a post-translational positive feedback loop whose activation is linked to a critical cell size (Tyson *et al*., 2002; Gerard *et al*., 2015).

In the absence of mitotic cyclin (e.g., cells carrying the *cdc13*^+^ gene driven by a thiamine-repressible promoter), fission yeast cells cannot enter into mitosis, but their Cdk1:Cig2 activity continues to oscillate (Wuarin *et al*., 2002). In response to these persisting Cdk1:Cig2 oscillations, cells execute periodic DNA replication without any mitosis and cell division (Hayles *et al*., 1994). During these ‘endoreplication’ cycles, the activation of Cig2-dependent kinase (‘Cig2-kinase’, hereafter) maintains its correlation to cell size, as indicated by a characteristic nucleocytoplasmic ratio at initiation of periodic DNA replications (Kiang *et al*., 2009).

DNA replication is also uncoupled from mitosis in *cdc18*^op^ cells that overexpress Cdc18 protein, a licensing factor for DNA replication (Nishitani and Nurse, 1995). Cdc18 is also an inhibitor of Cdc13-kinase. (Greenwood *et al*., 1998; Hermand and Nurse, 2007). Consequently, in cells overexpressing Cdc18, DNA origins of replication are constitutively licensed, and they have sufficient Cdk1 activity to initiate and reinitiate DNA replication but not enough to enter mitosis. These cells ‘over-replicate’ their genome in a temporally unorganized fashion. We also discuss situations when Cdc13-kinase activity is driven to low level by overexpression of Ste9, a component of the Cdc13 degradation pathway (Yamaguchi *et al*., 1997; Kitamura *et al*., 1998), or by overexpression of Rum1, a strong stoichiometric inhibitor of Cdk1:Cdc13 (Moreno and Nurse, 1994).

In this paper, we present a dynamical model of the Cdk1:Cig2 oscillations regulating fission-yeast DNA replication with special emphasis on endoreplication cycles. We build our model in a modular fashion starting with the transcriptional negative feedback loop, which is at the heart of the oscillatory mechanism. Nonlinear negative feedback is necessary, but not sufficient, for limit cycle oscillations, because the feedback signal must be ‘delayed’ in time (Novak and Tyson, 2008). The ‘delay’ could be an explicit time delay caused, e.g., by a long feedback loop, or it could be a functional delay caused, e.g., by positive feedback loop(s) that amplify the action of the activator or the inhibitor in the negative feedback loop (Novak and Tyson, 2008). Following this line of thinking, we propose two potential positive feedback mechanisms—not mutually exclusive—that could provide the necessary time-delay to make the negative feedback oscillatory. Next, we propose mechanisms for the entrainment of Cig2 oscillations by the nucleocytoplasmic ratio during endoreplication cycles and by Cdk1:Cdc13 oscillations during mitotic cycles. Finally, we discuss the molecular mechanisms that account for the difference between aperiodic DNA over-replication in some mutant strains and periodic endoreplication in other strains.

## Results

### The core negative feedback loop

Eukaryotic DNA replication origins must be ‘licensed’ before they become activated by Cdk1:cyclin complexes at initiation. In fission yeast, the role of DNA-replication licensing factor is played by Cdc18 (Kelly *et al*., 1993) and Cdt1 (Nishitani *et al*., 2000). The transcription of both licensing factors and the S-phase cyclin, Cig2, is induced by a transcription factor called MBF (MluI cell cycle Box-binding Factor), which contains the Cdc10 protein as a core component (Reymond *et al*., 1993). The activity of MBF fluctuates during both mitotic (Rustici *et al*., 2004) and endoreplication cycles (Kiang *et al*., 2009), being activated before and inactivated after DNA replication. The oscillation of MBF activity is caused by a transcriptional negative feedback loop created by several protein products of MBF-dependent transcription: Cig2, Nrm1 and Yox1. The Nrm1:Yox1 complex is thought to be a stoichiometric inhibitor of MBF (Cdc10) after S phase (Bertoli *et al*., 2013), creating a two-component negative feedback loop (Cdc10 ⟶ Nrm1 **—**| Cdc10; Figure 1B). In addition, Nrm1 protein is targeted for degradation by APC/C^Ste9^ (Bertoli *et al*., 2013), and Cdk1:Cig2 kinase (Ayte *et al*., 2001) inactivates APC/C^Ste9^ by multisite phosphorylation (Blanco *et al*., 2000; Yamaguchi *et al*., 2000), creating a longer negative feedback loop (Cdc10 ⟶ Cig2 **—**| Ste9 **—**| Nrm1 **—**| Cdc10; Figure 1B). During endoreplication cycles, which are our interest here, Cdk1:Cdc13 level is very low (*cdc13*^+^ transcription is repressed); therefore, the phosphorylation-dependent inactivation of Ste9 relies exclusively on Cdk1:Cig2. The S-phase Cdk1 complex can inactivate Ste9 efficiently because Cig2 is not targeted for degradation by APC/C^Ste9^ (Blanco *et al*., 2000). Cdk1:Cig2 also phosphorylates Cdc18, which targets the licensing factor for rapid ubiquitinylation-dependent degradation (Jallepalli *et al*., 1997). (In the following, we focus on Cdc18 regulation, neglecting Cdt1.) Notice that, in our model in Figure 1B, Cig2-kinase indirectly inhibits Cdc10 by inhibiting degradation of Nrm1 through inactivation of Ste9. In summary, the level of Nrm1 is upregulated by MBF-dependent transcription through a coherent feedforward loop that increases Nrm1 synthesis (Cdc10 ⟶ Nrm1) and inhibits its degradation (Cdc10 ⟶ Cig2 **—**| Ste9 **—**| Nrm1).

The inactivation of Ste9 by Cdk1:Cig2-dependent multisite phosphorylation and the stoichiometric inhibition of Cdc10 by Nrm1:Yox1 introduce strong nonlinearities into the transcriptional negative feedback loop. These nonlinear interactions can be conveniently illustrated by balance curves on an [Nrm1]_T_ – [Cig2] coordinate system, which is called a ‘phase plane’ (Figure 1C) (Novak and Tyson, 2008). A balance curve indicates the effect of one variable on the steady state level of the other. The balance curve of Cig2 (the green curve) decreases with increasing Nrm1, because Nrm1 inhibits Cdc10-dependent transcription of Cig2. In contrast, the steady state level of Nrm1 shows a switch-like increase above a threshold level of Cig2, the level at which Cig2-kinase effectively inactivates APC/C^Ste9^.

A negative feedback loop with sufficient nonlinearities and time-delay can exhibit sustained oscillation over a particular range of kinetic parameters (Novak and Tyson, 2008). The longer negative feedback loop in Figure 1B has four components, but its oscillatory potential is suppressed because the phosphorylation of Ste9 by Cig2-kinase and the inhibition of Cdc10 by Nrm1-binding are fast reactions compared to synthesis and degradation of Cig2 and Nrm1 proteins. By making pseudo-steady state assumptions on Ste9 and Cdc10, we reduce the four-dimensional dynamical system to two dimensions (the Cig2-Nrm1 phase plane in Figure 1C). The steady state of the reduced system is found at the intersection of the two balance curves, where both Cig2 and Nrm1 are both unchanging in time. This steady state is globally stable. According to numerical simulations (not shown), all components approach their steady state values monotonically, except Cdc18, which shows a transient peak because Cdc18 level is regulated by an incoherent feedforward loop: Cdc10 directly activates synthesis of Cdc18 and indirectly promotes its degradation (through Cig2-kinase). We note here that proteasomal degradation of Cdc18 and Cig2 is initiated by polyubiquitinylation of the phosphorylated substrates (Cdc18-P, Cig2-P) by the Skp-Cullin-Fbox (SCF) ubiquitin ligase (Toda *et al*., 1999).

Because the time delays in our transcriptional negative feedback loops (Cdc10 ⟶ Nrm1 **—**| Cdc10 and Cdc10 ⟶ Cig2 **—**| Ste9 **—**| Nrm1 **—**| Cdc10) are not long enough to induce oscillations, we consider positive feedback loops as alternatives to destabilize the steady state and generate relaxation-type oscillatory behaviour in the model. Since the transcriptional network has potential positive feedback loops, we explored these possibilities further.

### Positive feedback regulation of Cig2 activity

Cdk1:cyclin complexes are regulated by transient, inhibitory phosphorylations of their Cdk1 subunit which supresses their protein kinase activities. In fission yeast, the inhibitory Cdk1-phosphorylations provide the dominating control over mitotic Cdk1:Cdc13 complexes (Nurse, 1990). Moreover, the S-phase inducing Cdk1:Cig2 complex also undergoes transient, inhibitory phosphorylations, which persist if DNA replication is inhibited (Zarzov *et al*., 2002; Hermand and Nurse, 2007). In fission yeast, two inhibitory kinases, Wee1 (Russell and Nurse, 1987) and Mik1 (Lundgren *et al*., 1991), and two activatory protein-phosphatases (Cdc25 and Pyp3) are responsible for reversible, inhibitory phosphorylations of Cdk1:cyclin complexes (Russell and Nurse, 1986; Millar *et al*., 1992). The phenotypes of loss-of-function mutants suggest that Wee1 and Mik1 are acting, respectively, in late G2 phase and in S phase of the cell cycle (Nurse, 1975; Lundgren *et al*., 1991). Based on these observation, we propose that Mik1 is the dominant inhibitory kinase for the Cdk:Cig2 complex, which is a Cdc10 regulated protein (Ng *et al*., 2001). For simplicity, we assume constant activity for the activatory phosphatase (Cdc25 and/or Pyp3). The level of Mik1 protein fluctuates during the cell cycle, showing a peak during S phase (Christensen *et al*., 2000), which we explain by SCF-induced degradation of Mik1 after it is phosphorylated by Cig2-kinase, similar to Cdc18 degradation. Briefly, in our model, Cdk1:Cig2 and Mik1 are mutually inhibiting each other, which creates a positive feedback loop (Figure 2A).

**Figure 2:**
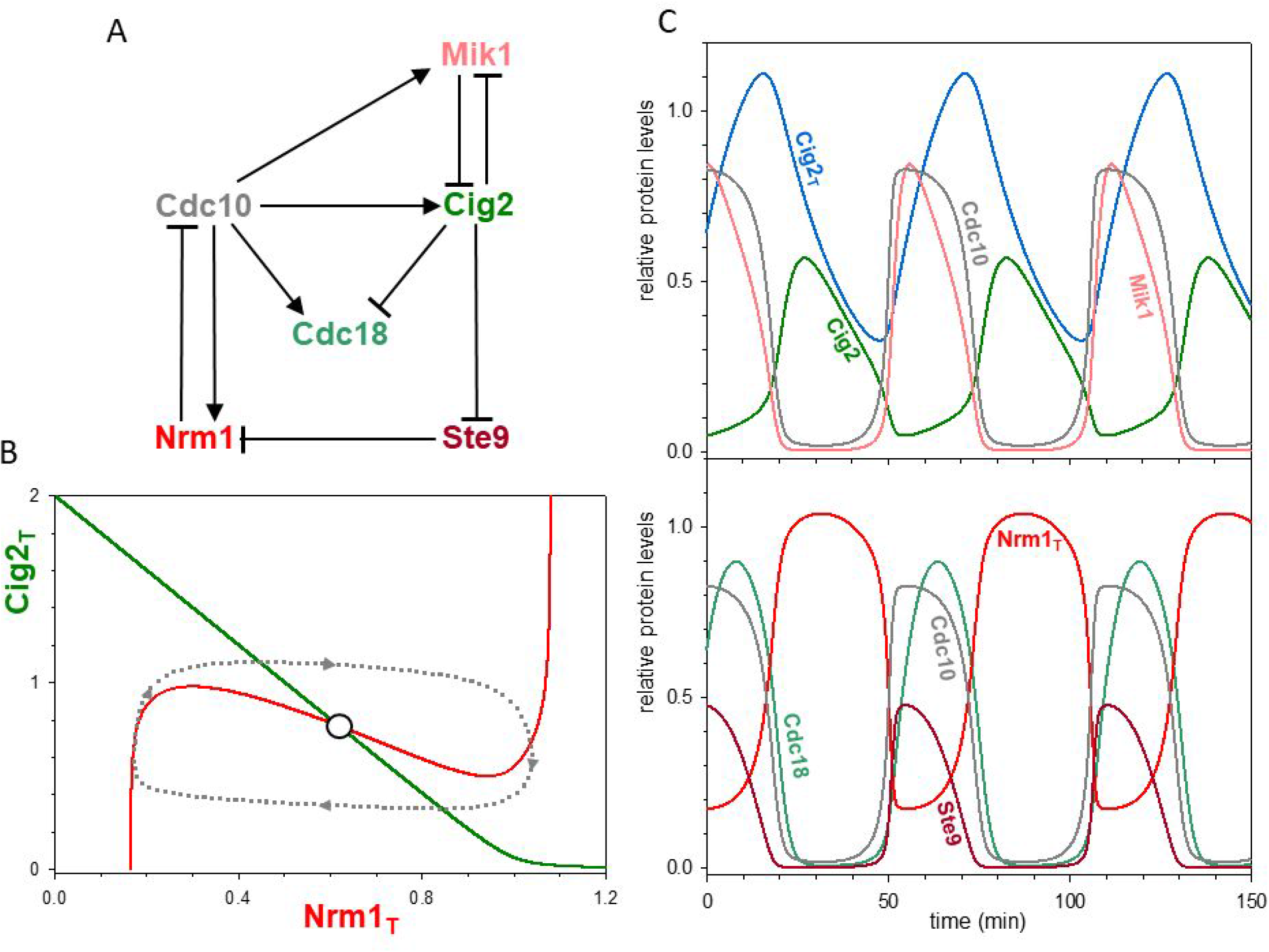
Positive feedback regulation of Cig2-kinase. (A) Influence diagram. (B) Phase plane portrait: the positive feedback makes Nrm1_T_ a bistable switch indicated by its N-shaped balance curve. The limit cycle oscillation is shown by the dotted closed orbit. (C) Numerical simulation of the network shown in (A).

The positive feedback loop puts a ‘kink’ in the Nrm1-balance curve (the red curve in Figure 2B) and thereby creates the possibility of oscillations in the Cig2-Nrm1 negative feedback loop (Novak and Tyson, 2008). The Nrm1-balance curve is N-shaped because more Cig2 is required to turn off Ste9 and upregulate Nrm1 than is needed to maintain Nrm1 at high levels. Notice that we plot on the y-axis the <u>total</u> Cdk1:Cig2 level, [Cig2]_T_, which is the sum of active and inactive (Cdk1 phosphorylated) complexes. This antagonistic relationship between Mik1 and Cdk1:Cig2 generates an effective time-delay to make the transcriptional negative feedback loop oscillatory (Figure 2C). During the oscillation, Mik1 inactivates Cdk1:Cig2 first, and only later is Cdk1:Cig2 able to promote Mik1 degradation. The delayed activation of Cdk1:Cig2 by inhibitory phosphorylation could be very significant for the control of DNA replication by providing a time-window for Cdc18 to licence replication origins before Cdk1:Cig2 activity appears (Figure 2C).

### Positive feedback regulation of Cdc10 transcription factor

In the previous example, the transcriptional negative feedback loop was made oscillatory by positive feedback regulation of Cig2 activity. The other possibility for introducing positive feedback is auto-activation of the Cdc10 transcription factor mediated by a ‘transcription factor inhibitor’ (TFI) (Figure 3A). The paradigm for this hypothetical TFI is the regulation of transcription factors E2F and SBF in mammalian cells and budding yeast, respectively (see e.g. (Bertoli *et al*., 2013)). Both E2F and SBF are inhibited by stoichiometric binding partners (Retinoblastoma protein and Whi5, respectively) which become inactivated by Cdk:cyclin complexes (Cdk2:CycE and Cdk1:Cln2, respectively) that are upregulated by their respective transcription factors. Since fission yeast also has a Whi5 gene and protein, it is reasonable to hypothesize a stoichiometric inhibitor (TFI) of Cdc10 that controls the activity of MBF. We assume that unphosphorylated TFI binds reversibly and inhibits MBF (Cdc10) and the inhibition is terminated by phosphorylation of TFI, which is carried out by a protein kinase upregulated by Cdc10. Since the Cdc10 target responsible for inactivation of TFI is unknown, we assume for the time being that Cig2 is responsible for this reaction.

**Figure 3:**
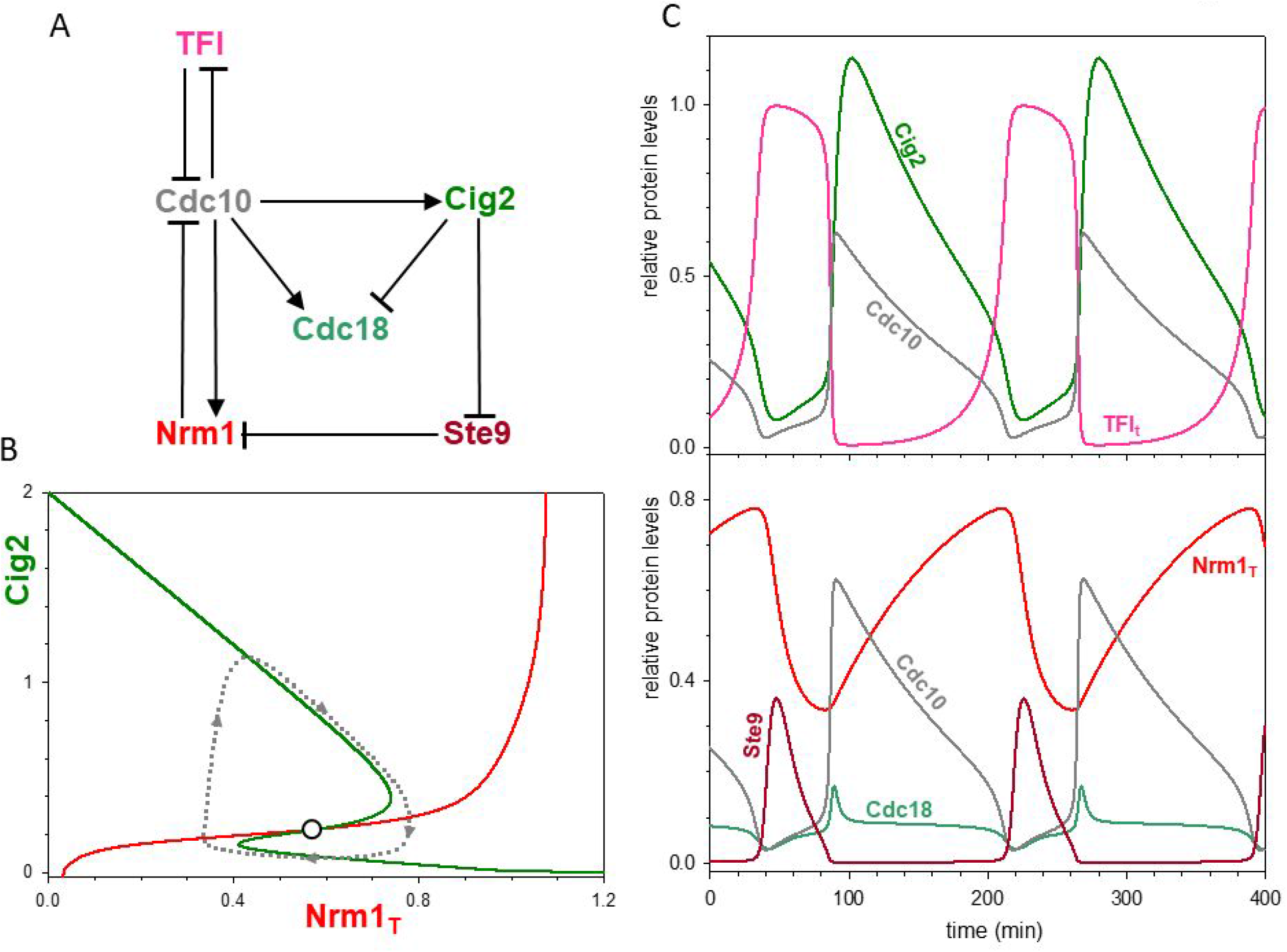
Positive feedback regulation of Cdc10 transcription factor by a G1 specific inhibitor (TFI). (A) Influence diagram. (B) Phase plane portrait: the Z-shaped balance curve indicates that Cig2-kinase behaves as a bistable switch. The limit cycle oscillation is shown by the dotted closed orbit. (C) Numerical simulation of the network shown in (A).

In this scenario, MBF has two stoichiometric inhibitors, which are regulated differently: activation of Cdc10 inhibits TFI while it activates Nrm1 directly (synthesis) and indirectly through Cig2 and Ste9. The positive feedback loop between Cdc10 and TFI puts a kink in the Cig2 balance curve (the green curve in Figure 3B). On the upper and the lower branches of the Cig2 balance curve TFI is phosphorylated (inactive) and dephosphorylated (active), respectively. The bistable region is bounded by two different Nrm1 thresholds. The Nrm1 threshold required to induce dephosphorylation of TFI is higher than the threshold required to maintain TFI in the active, dephosphorylated state. If the red balance curve intersects the Z-shaped green balance curve on the middle branch of the ‘Z’, then the system exhibits limit cycle oscillations with periodic phosphorylation and dephosphorylation of TFI (Figure 3C). The oscillations of Cig2, Nrm1 and Cdc10 are quite acceptable in this model, but the waveform of Cdc18 oscillations is unacceptable. Cdc18 (the licensing factor) cannot accumulate significantly because there is little time-delay between activation of Cdc10 and of Cig2.

The two positive feedback mechanisms controlling Cdc10 and Cig2 activities are not mutually exclusive; most likely, they operate in parallel in the same regulatory network (Figure 4A). With both positive feedbacks in place, both Nrm1 and Cig2 balance curves are kinked (N-and Z-shaped, respectively, in Figure4B), and the oscillations preserve the good dynamic features of both scenarios (Figure 4C). Although the positive feedback on Cdc10 seems to be inconsequential in the oscillation, it plays an important physiological role as we discuss in the next session.

**Figure 4:**
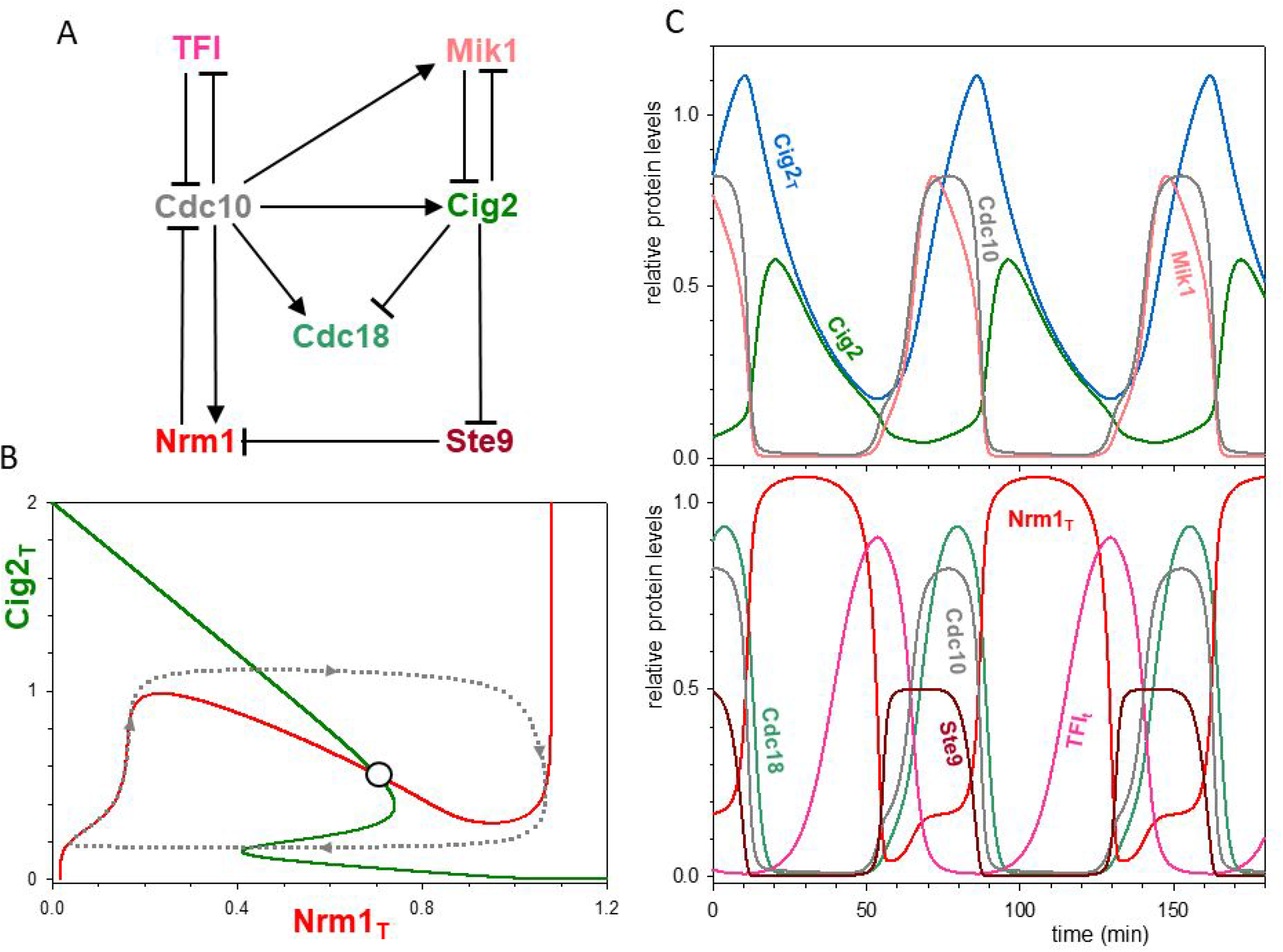
The transcriptional negative feedback regulated by two positive feedback loops. (A) Influence diagram. (B) Phase plane portrait: both Nrm1_T_ and Cig2_T_ dis[lay bistability, as indicated by their N-and Z-shaped balance curve, respectively. The limit cycle oscillation is shown by the dotted closed orbit. (C) Numerical simulation of the network shown in (A).

### Control by the nucleocytoplasmic ratio

During endoreplication, Cig2-kinase oscillations drive discrete rounds of DNA synthesis with the periodicity of cell mass doubling time (Kiang *et al*., 2009), which suggests a strong coupling between the oscillatory mechanism in Figure 4 and the nucleocytoplasmic ratio (NCR). The NCR oscillates because the cellular DNA content doubles sharply during DNA replication and declines slowly by cell growth outside S phase. It is reasonable to assume that endoreplication (when cells skip mitosis) is controlled by the NCR at the G1/S transition when Cdc10 is activated by relieving it from inhibition by TFI. In general, the period of the Cig2-kinase oscillation becomes adjusted to the cell mass doubling time if a component in the Cdc10 transcriptional network mirrors the NCR. One possibility is an ‘inhibitor-dilution mechanism’ (Fantes *et al*., 1975), for which the NCR is mirrored by the intracellular concentration of TFI. This mechanism requires that the TFI synthesis rate is dictated by gene-dosage rather than by the size of the cell (Schmoller *et al*., 2015; Heldt *et al*., 2018).

An inhibitor-dilution model is explored in Figure S1. Figure S1A illustrates how the total concentration of TFI affects the dynamic properties of the network. The network can only oscillate at low TFI concentrations; when [TFI]_T_ is higher than a threshold level (the red arrow in Figure S1A), the network admits only a stable G1 steady-state (‘G1 arrest’), with inactive Cdc10 transcription factor. The stable steady state is created near the origin of the phase plane (low Cig2 and Nrm1 levels) by the intersection of the two balance curves (Figure S1B). In this model (constant TFI synthesis rate during growth), [TFI]_T_ is steadily decreasing as the cell grows, and the cell’s dynamic state is slowly drifting to the left on Figure S1A. Once [TFI]_T_ drops below the threshold of the stable steady-state regime, the network starts to oscillate by activation of Cdc10. The rise of Cdk1:Cig2 activity initiates DNA replication, which results in duplication of the TFI gene and a doubling of its rate of synthesis. This will lead to an increase of total TFI concentration (assuming fast TFI turn-over), which pushes the dynamic state of the network back above the star, where it becomes attracted to the stable G1 steady-state again. Numerical simulations show that this TFI-dilution mechanism only results in NCR-controlled endoreplication cycles if the TFI level rapidly mirrors changes of nucleocytoplasmic ratio (simulations not shown). However, rapid change of TFI concentration requires that the TFI protein is unstable, which does not seem to be the case for budding yeast Whi5, which has a half-life longer than 6 hrs (Schmoller *et al*., 2015).

We rather favour a ‘titration-mechanism’ for which the number of a starter kinase (SK) molecules per genome mirrors the inverse of NCR (Figure 5A). We envision a SK molecule that binds strongly to specific binding sites on chromosomes where it inactivates TFI in complex with Cdc10 by phosphorylation. SK molecules accumulate in an amount proportional to cytoplasmic contents (e.g., ribosomes) and bind to sites proportional to nuclear chromosome number. This mechanism makes the rate of TFI inactivation by SK proportional to the inverse of NCR and it leads to Cdc10 activation above a critical inverse NCR value. One of the kinases upregulated by Cdc10 also phosphorylates and inactivates TFI, and this positive feedback loop causes an abrupt entry into S phase. Figure 5B illustrates how the total concentration of SK affects the dynamic properties of the network. At low [SK], there exists only a stable steady stat of low Cig2-kinase activity (G1 arrest). As the cell grows, [SK] eventually crosses a threshold (the red arrow in Figure 5B), past which the G1 arrested state disappears and the control system sets off on an oscillation. As before, rising Cig2-kinase activity initiates DNA replication, which doubles the number of chromosomal binding sites and halves the number of SK molecules per genome, pushing the control system back into the domain of attraction of the stable G1 steady state. Numerical simulation of this titration-mechanism provides a robust NCR-controlled endoreplication cycle (Figure 5C), where the temporal period of DNA replication is identical to the mass doubling time of the cells.

**Figure 5:**
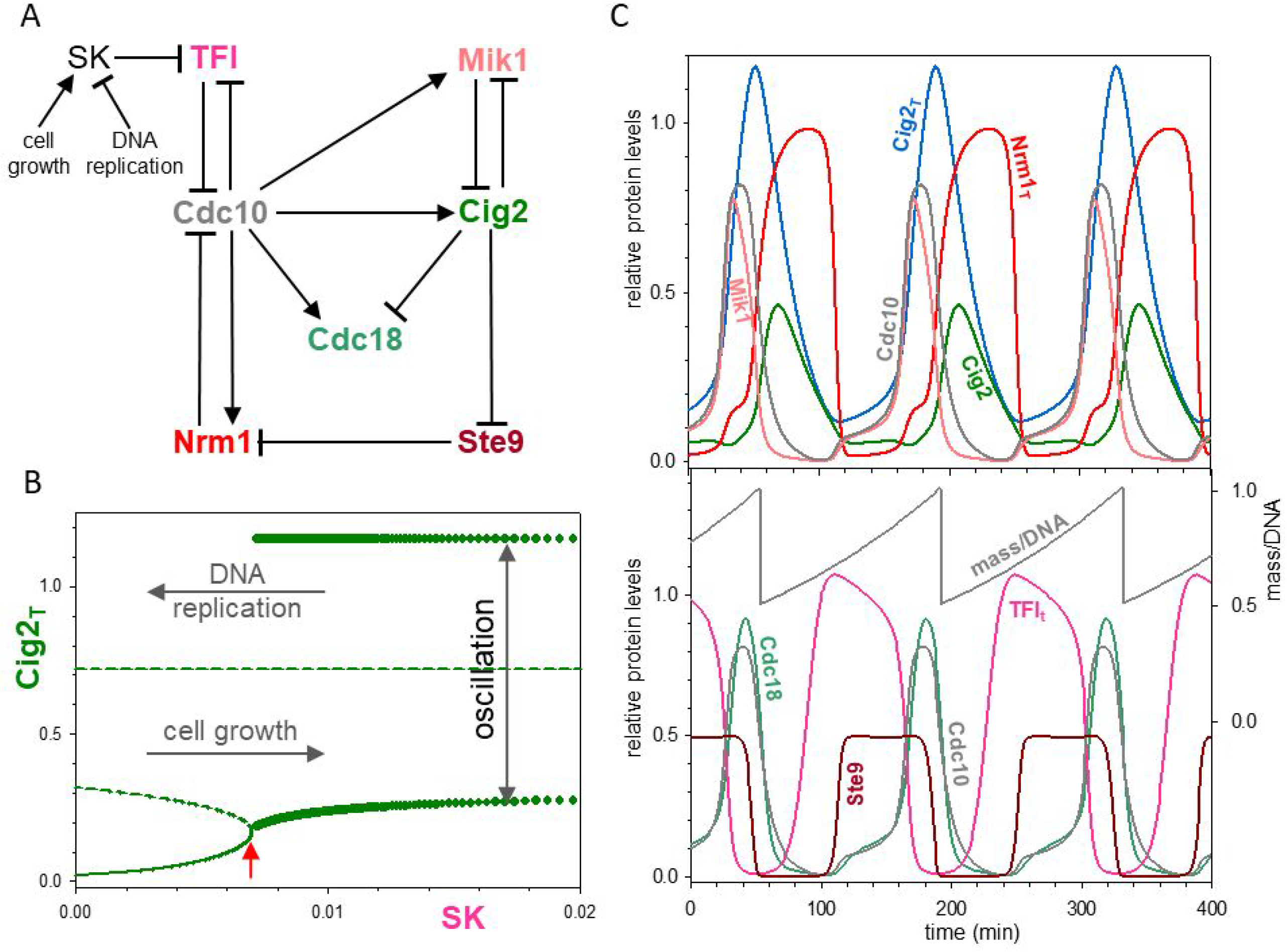
Nucleocytoplasmic control of the oscillation. (A) Influence diagram of the network with a starter kinase (SK) whose activity is controlled by nucleocytoplasmic ratio (NCR). (B) The effect of SK level on the regulatory network. Small SK levels create stable steady states (solid line), but high levels of SK allow large amplitude oscillations (the filled circles are minima and maxima). Unstable steady states indicated by dashed lines. (C) Numerical simulation of the network.

### The DNA-replication checkpoint

In the previous section, we showed how oscillations in the nucleocytoplasmic ratio (NCR) can entrain the transcriptional oscillator to the mass doubling time of cell growth. High values of the NCR stabilize a G1-arrested state, i.e. a stable steady state of low activity of Cdc10 transcription factor and Cig2-kinase activity. In this section we discuss the effect of a ‘replication checkpoint’ on the Cig2-kinase oscillation. Inhibition of DNA replication by hydroxyurea leads to stabilization of both total Cig2 level and the phosphorylated (inhibited) form of Cdk1:Cig2 (Zarzov *et al*., 2002; Hermand and Nurse, 2007), as well as dissociation of Nrm1 from MBF (de Bruin *et al*., 2008). Experiments also suggest that Cdc18 (licensing factor) is necessary for maintenance of inhibitory phosphorylation of Cdk1 and thereby for the replication checkpoint (Hermand and Nurse, 2007). We implement these experimental findings in our model (see Figure 6A) by supposing that unreplicated DNA (1) inhibits Cig2-kinase induced degradation of Mik1 in a Cdc18-dependent manner, and (2) increases the equilibrium dissociation constant between Cdc10 and Nrm1. These parameter changes influence both balance curves in the following ways (see Figure 6B). The [Cig2]_T_ balance curve (green) becomes horizontal, because Cdc10 inhibition by Nrm1 is compromised. The N-shaped characteristic of the Nrm1 balance curve (red) is lost because of the weaker antagonism between Cig2-kinase and Mik1. These changes of the balance curves create a new intersection point with high Cig2 level (but low Cig2-kinase activity) and intermediate Nrm1 level, which defines a stable steady state corresponding to the replication checkpoint.

**Figure 6:**
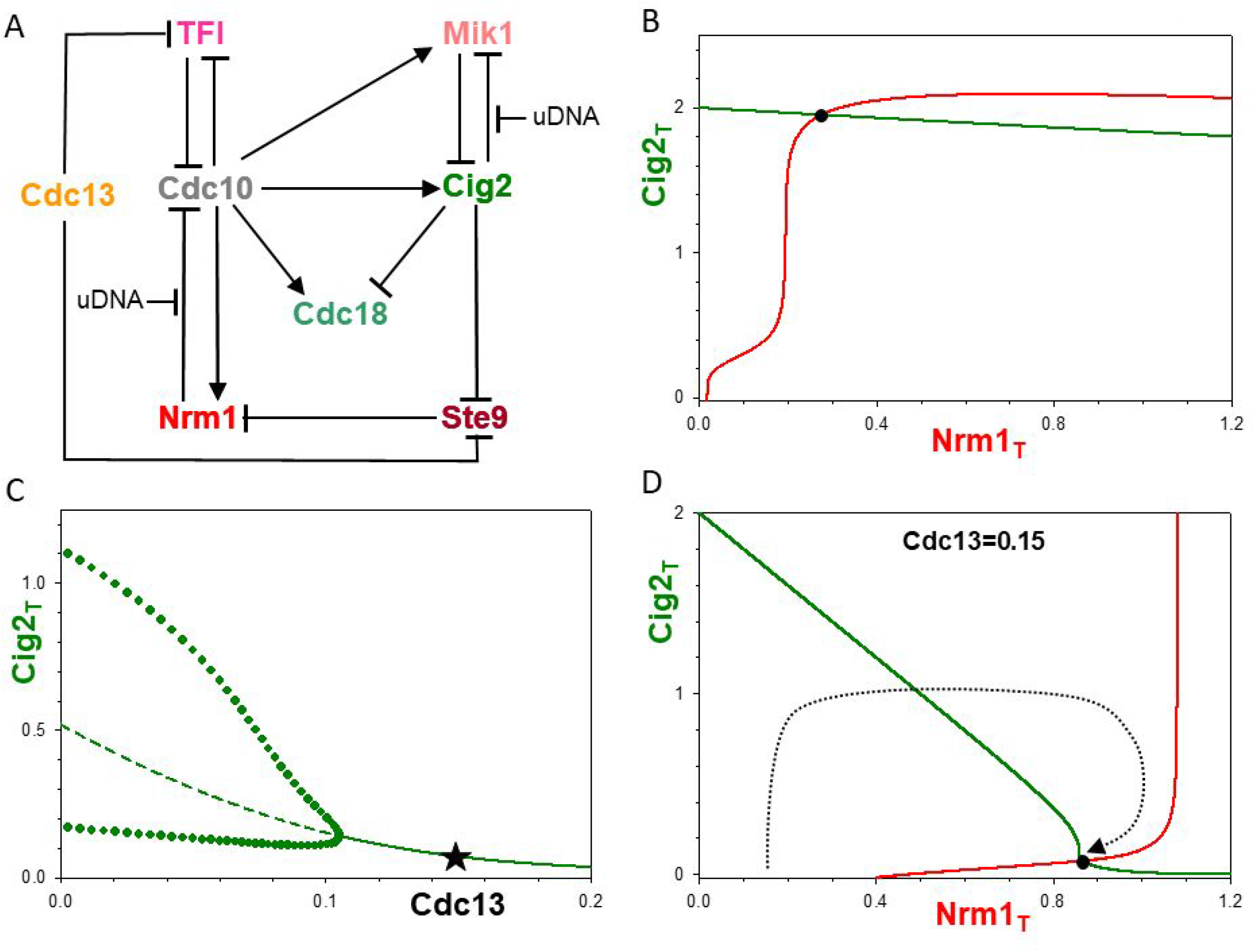
The effect of replication checkpoint and Cdc13 mitotic cyclin on the oscillation. (A) The effect of Cdc13-kinase and unreplicated DNA on the network. (B) Unreplicated DNA creates a stable steady state (●) corresponding to the replication checkpoint. (C) The effect of Cdc13-kinase activity on the oscillation. The gradual decrease of the oscillation amplitude (indicated by filled circles) terminates in a stable steady state (solid line) by a Hopf-bifurcation. (D) Higher than threshold level of Cdc13 creates a stable steady state corresponding to a G2-like state (Cig2 level lo, Nrm1 high).

### Entraining Cig2-kinase activity to Cdc13-kinase oscillations

Another important question concerns the mechanism that couples the Cdk1:Cig2 and Cdk1:Cdc13 oscillators during mitotic cycles. The strict alternation of Cdk activities driving S-phase and M-phase of the mitotic cycle (Figure 1A) requires suppression of the Cdk1:Cig2 oscillator by Cdk1:Cdc13 kinase during G2 and M phases. The overlapping substrate specificity of Cdk1:cyclin complexes provides a simple solution for tight coupling of the two oscillators. By making both Ste9 and TFI phosphorylable substrates of Cdk1:Cdc13, the Cdk1:Cig2 oscillator becomes sensitively dependent on the <u>absence</u> of the mitotic kinase (Figure 6C), consistent with fundamental observations by Hayles et al. (Hayles *et al*., 1994). With increasing Cdk1:Cdc13 activity, the large amplitude Cig2-oscillations are damped down to a stable steady state (Figure 6C). This steady state, created by Cdk1:Cdc13 phosphorylation of Ste9, is located in the bottom-right corner of the Nrm1-Cig2 phase plane (Figure 6D), with high Nrm1 and low Cig2 levels that are characteristic of G2 cells. The stable steady state is created by flattening the Nrm1 balance curve, a consequence of Cdc13-dependent inactivation of Ste9, which stabilizes Nrm1. In the presence of Cdc13 (mitotic cyclin), the Cig2-oscillator cannot complete a full cycle, because the stable steady state in Figure 6D represents a G2-M ‘checkpoint’. Only after mitotic degradation of Cdc13 will this checkpoint be removed, allowing the Cig2-oscillator to proceed into a G1-like state. These simulations demonstrate that our model of endoreplication can also explain the strict alternation of S and M phases during mitotic cycles.

### Uncoupling of DNA replication from mitosis in other situations

In addition to depletion of Cdc13 mitotic cyclin, multiple rounds of DNA replication in the absence of mitosis can be induced by overexpression of several cell cycle regulators (*cdc18*^op^, *rum1*^op^ or *ste9*^op^) in *cdc13*^+^ genetic background (Moreno and Nurse, 1994; Yamaguchi *et al*., 1997; Kitamura *et al*., 1998). Rum1 is a strong stoichiometric inhibitor of Cdk1:Cdc13 complex (Correa-Bordes and Nurse, 1995); Ste9/Srw1 is an activator of APC/C-dependent degradation of Cdc13 (Yamaguchi *et al*., 1997; Kitamura *et al*., 1998); Cdc18 plays a role in the inhibition of Cdk1:cyclin by the unreplicated DNA checkpoint (Hermand and Nurse, 2007). Therefore, overexpression of any one of these three cell cycle regulators has the ability to eliminate or countermand Cdc13-kinase activity and create a situation similar to Cdc13-depletion. However, overexpression of these cell cycle regulators has pleiotropic effects because of their influence on other proteins in the network, and these effects must be taken into consideration in the model. While Ste9 has a direct pleiotropic effect in our network by promoting Nrm1 degradation, both Ste9 and Rum1 have indirect effects by weak inhibition of Cdk1:Cig2 kinase (Correa-Bordes and Nurse, 1995; Yamaguchi *et al*., 1997; Benito *et al*., 1998). Furthermore, very high expression of Cdc18 causes strong inhibition of Cdk1:Cdc13 activity because excessive binding of Cdc18 to the kinase out-competes other Cdk1 substrates (Greenwood *et al*., 1998).

The effect of overexpression of these cell cycle regulators can be conveniently illustrated on a Signal-Response (S-R) diagram where the Signal is the extent of protein overexpression and the Response is one of the network components (we choose the total Cig2 level). The S-R diagrams for overexpression of Cdc18, Rum1 or Ste9, shown in Figure S2, are similar because the effects of overexpression on the Nrm1-balance curve (the red curves in Figure7) are similar, as we discuss below. Referring again to Figure S2, we see that, at low expression levels of these proteins, the Cig2-oscillator is arrested in a G2/M-like state, as usual during mitotic cycle, by inhibition of Ste9 by the mitotic Cdc13-kinase. This stable steady state, with low Cig2 level, is only observable during mitotic cycles, and its elimination is required for uncoupling DNA replication from mitosis. In the middle regime of the bifurcation diagrams in Figure S2, the two chromosomal events (S and M) become uncoupled, and the Cig2-control network shows limit cycle oscillations similar to Cdc13-depleted cells. In the oscillatory regime, the two balance curves of the phaseplane (Figure7C,D) intersect on the middle branch of the N-shaped Nrm1-balance curve in an unstable steady state. Despite similar phase plane portraits and S-R diagrams, the temporal dynamics of *ste9*^op^ and *rum1*^op^ oscillations are quite different (Figure S3). The oscillations in *ste9*^op^ are indistinguishable from the cdc13-depleted situation (Figure5). On the other hand, overexpression of Rum1 significantly inhibits Cig2-kinase activity, thereby delaying DNA replication and accumulation of Nrm1 (Rum1 **—**|Cig2 **—**| Ste9 **—**| Nrm1). This delay of the onset of DNA replication causes a significant (∼40%) increase of cell size-to-DNA ratio in *rum1*^op^ cells compared to *cdc13*-depleted cells, in agreement with experimental data (Moreno *et al*., 1994; Kiang *et al*., 2009).

**Figure 7:**
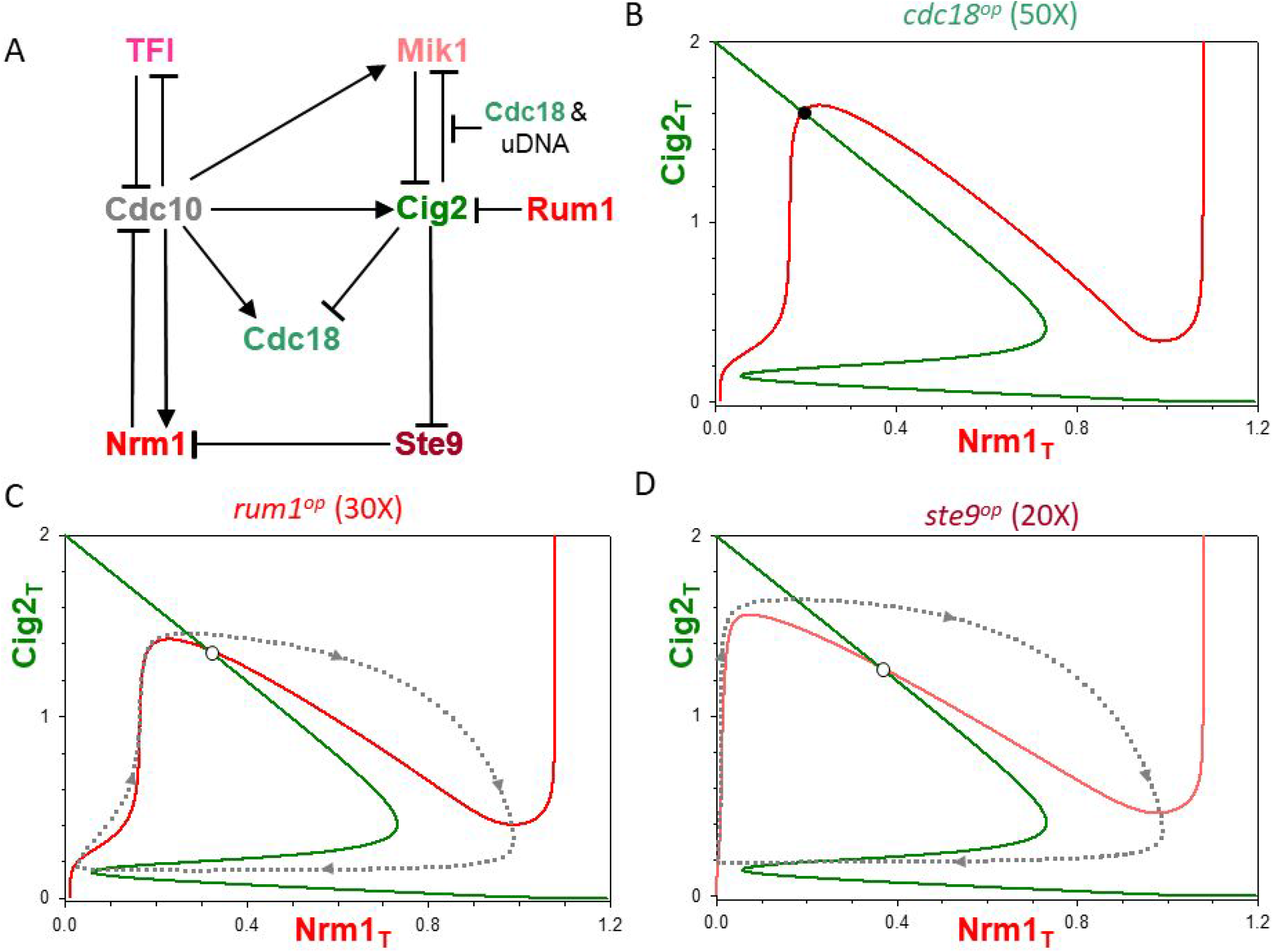
Re-replication in overexpressing strains. (A) Influence diagram. Phase plane portraits for overexpression of *cdc18* (B), *rum1* (C) and *ste9* (D). In all cases, overexpression of proteins increases the N-shaped Nrm1_T_ balance curve compared to a Cdc13-depleted strain. The levels of overexpression of Rum1 and Ste9 in panels C and D drive oscillatory endoreplication cycles (see Figure S3), corresponding to the oscillatory regimes of Figure S2. The *cdc18*^op^ level in panel B locks the network in a stable steady state with high-Cig2_T_ and low-Nrm1_T_ levels, which we associate with DNA over-replication.

In the oscillatory regime, the unstable steady state, at the intersection of the two balance curves on Figure7, is raised proportionally to the extent of overexpression of the modulated protein (Cdc18, Rum1, or Ste9). The increase of the unstable steady state is caused by the rise of the N-shaped Nrm1-balance curve (the red curve in Figure7), because high levels of these regulators of Cdk1 activity make it harder for Cdk1:Cig2 to inactivate Ste9 than in Cdc13-depleted cells. Eventually, the rise of the Nrm1-balance curve destroys the limit cycle oscillations, above critical levels of overexpression, thereby creating a third regime where the control system settles in stable steady state of high levels of Cig2, Cdc18 and Mik1 (Figure S2). Although the total level of Cig2 is high, the activity of Cdk1:Cig2 kinase is moderated by tyrosine phosphorylation of Cdk1 by Mik1. Cig2-kinase activity is below the threshold for Cdc10 inhibition, so the Cdc10-targeted genes (*cdc18, cig2* and *mik1*) are highly expressed. But the Cig2-kinase activity is high enough to support DNA replication in these mutant cells, as indicated by their phenotypes.

For example, the phaseplane on Figure 7B illustrates the intersection of the Cig2 and Nrm1 balance curves for a high level of Cdc18 overexpression, sufficient to create a stable steady state with high Cig2 and low Nrm1 levels. We associate this stable steady state with the phenotype characteristic for *cdc18*^*op*^ cells. Because their replication origins are simultaneously licensed and activated, Cdc18-overexpressing cells are continuously engaged in DNA replication, a phenotype called over-replication (Nishitani and Nurse, 1995). In this steady state, the protein levels of Cdc18, Cig2 and Mik1 are all upregulated, thereby creating a condition where both the licensing factor (Cdc18) and the activator (Cig2-kinase) of DNA replication coexist permanently. Although Cig2-kinase activity is depressed by Mik1, it is apparently large enough to drive disorganized DNA replication in these cells. Observe that the similarities of the steady states of Cdc18-overexpressing cells (Figure7B) and of S phase-arrested cells by HU (hydroxyurea); in both cases, Cig2 level is high and Nrm1 level is low. The difference is that HU blocks DNA synthesis by depletion of its precursors, while in *cdc18*^op^ cells, replication is ongoing.

According to our model, endoreplication transitions to over-replication with increasing levels of protein overexpression. Overexpression of gene transcription in fission yeast is accomplished using the thiamine-repressible *nmt* promoter (e.g., *nmt-rum1*^+^); overexpression is induced by removing thiamine from the growth medium. Since it takes many hours for fission yeast cells to metabolize residual, intracellular thiamine, the level of overexpression of the mutant gene rises steadily over the induction period. It is likely, therefore, that, as the protein level increases, there is a continuous shift of growing cells from mitotic cycles, to NCR-regulated endoreplication cycles, to continuous over-replicative genome expansion. Our model suggests the possibility of endoreplication for *cdc18*^op^ cells during early induction and over-replication for *rum1*^op^ and *ste9*^op^ cells at very high expression levels. These phenotypes have not yet been observed; they are predictions of the model that could be investigated experimentally.

## Discussion

During normal growth and proliferation, fission yeast cells exhibit strict alternation of DNA replication and mitotic division, in order to maintain their nucleocytoplasmic ratio (NCR = DNA/protein ratio) within strict bounds, an approximately two-fold range around an optimal ratio. The timing of DNA synthesis and mitosis during the fission yeast cell cycle is controlled by fluctuating activities of Cig2-dependent kinase and Cdc13-dependent kinase, respectively. Their oscillations are strongly coupled so that there is one peak of each kinase activity in each cell cycle; furthermore, the oscillations of DNA replication and cell division are strongly entrained to cell growth, so that wild-type fission yeast cells exhibit a stable and narrow distribution of cell length at division (14 ± 0.1 μm).

These properties of fission yeast cell reproduction can be compromised in mutant strains. If mitotic cyclin (Cdc13) level is sufficiently depleted, then nuclear and cell division are eliminated, but DNA replication continues by periodic genome doubling in concert with mass doubling, a phenomenon called NCR-controlled ‘endoreplication’. During endoreplication, origins of replication are periodically licensed (bound by Cdc18 licensing factor), fired (phosphorylated by Cig2-kinase activity), de-licensed (by degradation of Cdc18), and then re-licensed (by activation of Cdc10, a transcription factor for Cdc18 and Cig2). This sequence of events is periodically initiated by oscillations of Cdc10 transcription factor, as the mutant cells undergo endoreplication cycles: G1⟶S⟶G2⟶G1⟶ …. Overexpression of Cdc18 (replication licensing factor) in fission yeast induces over-replication of the genome without periodic execution of G1/S transitions (Nishitani and Nurse, 1995). In cells overexpressing Cdc18, DNA origins cannot be de-licensed, and high levels of Cig2 protein in these strains causes continuous, disorganized replication of chromosomes.

In this work, we have presented a new dynamic model of endoreplication cycles in fission yeast cells. At the heart of our model, the Cdc10 transcription factor is regulated by two negative feedback loops (NFLs, see Figure 1A). Cdc10 up-regulates synthesis of its own stoichiometric inhibitor, Nrm1; this is a short (two-component) NFL. Additionally, Cdc10 drives synthesis of Cig2, and Cig2-kinase inactivates an E3 ubiquitin ligase (APC/C^Ste9^) that labels Nrm1 for degradation by proteasomes (Figure 1A); this is a four-component NFL. Because neither of these NFLs can oscillate on its own, we supplement them with a positive feedback on Cig2-kinase activity by inhibitory phosphorylation of the kinase subunit (Cdk1, see Figure 2A). This addition creates an oscillation with temporal separation between DNA replication licensing (by Cdc18) and initiation (by Cig2). An additional (hypothetical, but likely) positive feedback controls the activation of Cdc10 by a transcription factor inhibitor (TFI, Figure 3A). This addition provides a means to couple the oscillator with cytoplasmic growth (Figure 5A). These NCR controlled oscillations drive multiple, discrete rounds of DNA replication (Figure 5) that we associate with endoreplication cycles observed in Cdc13-depleted fission yeast cells (Hayles *et al*., 1994).

In our earlier model for the endoreplication cycle (Novak and Tyson, 1997), Rum1, a G1 regulatory protein, played a central role in the oscillatory mechanism. Rum1 is stoichiometric inhibitor of Cdk1:cyclin complexes; a stronger inhibitor of Cdk1:Cdc13 than of Cdk1:Cig2. Rum1 degradation is promoted by Cdk:cyclin-dependent phosphorylation (Benito *et al*., 1998). In the early model, Cig2 level was regulated by a NFL, while its activity was amplified by the double-negative (i.e., ‘positive’) feedback between Rum1 and Cig2-kinase. In the present model, the role of Rum1 (to inhibit Cdk1:Cig2 kinase activity) is replaced Mik1 (and possibly by Wee1 as well), a protein kinase that phosphorylates Cdk1 and inhibits its kinase function. Rum1 is not part of the <u>core</u> Cig2 oscillatory mechanism driving endoreplication cycle (Figure 5A) in the present model. We propose that Rum1, although it regulates the activity of Cdk1:Cig2, is not necessary to drive Cig2 oscillations.

In addition to depletion of Cdc13 mitotic cyclin, multiple rounds of DNA replication in the absence of mitosis can be induced by overexpression of several cell regulators (*cdc18*^op^, *rum1*^op^ or *ste9*^op^) in *cdc13*^+^ genetic background. Interestingly, each of these proteins (Cdc18, Rum1 and Ste9), besides being a strong Cdk1:Cdc13 inhibitor, has significant pleiotropic effects on other components of the network, e.g inhibition of Cdk1:Cig2. As a result of such additional effects, our model suggests that mutants overexpressing these proteins do not necessarily correspond to the oscillatory behaviour (periodic endoreplication) characteristic for Cdc13-depleted cells; rather, sufficiently high levels of overexpression induce over-replication of the genome.

We explain the difference between endoreplication and over-replication in terms of our dynamical model using the concept of balance curves on a phase plane. Periodic endoreplication is understood to be a limit cycle oscillation on the Nrm1-Cig2 phase plane (Figure 4B) derived from an unstable steady state at the intersection of the Nrm1 (red) and Cig2 (green) balance curves. DNA over-replication in cells overexpressing Cdc18 suggests that *cdc18*^op^ cells are stuck in an unusual state where replication origins are simultaneously licensed and activated. In our model, constitutive overexpression of Cdc18 raises the N-shaped Nrm1-balance curve until a stable steady state is created in the upper left corner of the phase plane (Figure 7A). In this steady state, the protein levels of Cdc10-targeted genes (*cdc18, mik1* and *cig2*) are all upregulated. We associate this steady state (with high Cig2, <u>high Cdc18,</u> and low Nrm1 levels) to an over-replication phenotype. This observation leads us to conclude that endoreplication and over-replication are distinguished by a qualitative change of the dynamical system (a ‘bifurcation,’ in the language of dynamical systems theory) (Tyson and Novak, 2020).

## Materials and methods

All mathematical equations and computational methods are described and explained in detail in the Supplemental Text. We also provide the code for each model in the form of “.ode” files (see the Supplemental ODE Files) that allow the users to reproduce our figures with the free available software XPP/AUTO (http://www.math.pitt.edu/~bard/xpp/xpp.html).

## Acknowledgments

We are grateful to Jacky Hayles and Sergio Moreno for profitable discussions and to Adrien Wald for his contribution at an early stage of the project, We acknowledge financial support from BBSRC Strategic LoLa grant BB/M00354X/1 to B.N.

## Notes

### Competing Interest Statement

The authors have declared no competing interest.

## References

Ayte, J., Schweitzer, C., Zarzov, P., Nurse, P., and DeCaprio, J.A. (2001). Feedback regulation of the MBF transcription factor by cyclin Cig2. Nat Cell Biol 3, 1043–1050.

Benito, J., Martin-Castellanos, C., and Moreno, S. (1998). Regulation of the G1 phase of the cell cycle by periodic stabilization and degradation of the p25rum1 CDK inhibitor. EMBO J 17, 482–497.

Bertoli, C., Skotheim, J.M., and de Bruin, R.A. (2013). Control of cell cycle transcription during G1 and S phases. Nat Rev Mol Cell Biol 14, 518–528.

Blanco, M.A., Sanchez-Diaz, A., de Prada, J.M., and Moreno, S. (2000). APC(ste9/srw1) promotes degradation of mitotic cyclins in G(1) and is inhibited by cdc2 phosphorylation. EMBO J 19, 3945–3955.

Christensen, P.U., Bentley, N.J., Martinho, R.G., Nielsen, O., and Carr, A.M. (2000). Mik1 levels accumulate in S phase and may mediate an intrinsic link between S phase and mitosis. Proc Natl Acad Sci U S A 97, 2579–2584.

Correa-Bordes, J., and Nurse, P. (1995). p25rum1 orders S phase and mitosis by acting as an inhibitor of the p34cdc2 mitotic kinase. Cell 83, 1001–1009.

Coudreuse, D., and Nurse, P. (2010). Driving the cell cycle with a minimal CDK control network. Nature 468, 1074–1079.

de Bruin, R.A., Kalashnikova, T.I., Aslanian, A., Wohlschlegel, J., Chahwan, C., Yates, J.R., 3rd, Russell, P., and Wittenberg, C. (2008). DNA replication checkpoint promotes G1-S transcription by inactivating the MBF repressor Nrm1. Proc Natl Acad Sci U S A 105, 11230–11235.

Fantes, P.A., Grant, W.D., Pritchard, R.H., Sudbery, P.E., and Wheals, A.E. (1975). The regulation of cell size and the control of mitosis. J Theor Biol 50, 213–244.

Fisher, D.L., and Nurse, P. (1996). A single fission yeast mitotic cyclin B p34cdc2 kinase promotes both S-phase and mitosis in the absence of G1 cyclins. EMBO J 15, 850–860.

Gerard, C., Tyson, J.J., Coudreuse, D., and Novak, B. (2015). Cell cycle control by a minimal Cdk network. PLoS Comput Biol 11, e1004056.

Greenwood, E., Nishitani, H., and Nurse, P. (1998). Cdc18p can block mitosis by two independent mechanisms. J Cell Sci 111 (Pt 20), 3101–3108.

Hayles, J., Fisher, D., Woollard, A., and Nurse, P. (1994). Temporal order of S phase and mitosis in fission yeast is determined by the state of the p34cdc2-mitotic B cyclin complex. Cell 78, 813–822.

Heldt, F.S., Lunstone, R., Tyson, J.J., and Novak, B. (2018). Dilution and titration of cell-cycle regulators may control cell size in budding yeast. PLoS Comput Biol 14, e1006548.

Hermand, D., and Nurse, P. (2007). Cdc18 enforces long-term maintenance of the S phase checkpoint by anchoring the Rad3-Rad26 complex to chromatin. Mol Cell 26, 553–563.

Jallepalli, P.V., Brown, G.W., Muzi-Falconi, M., Tien, D., and Kelly, T.J. (1997). Regulation of the replication initiator protein p65cdc18 by CDK phosphorylation. Genes Dev 11, 2767–2779.

Kelly, T.J., Martin, G.S., Forsburg, S.L., Stephen, R.J., Russo, A., and Nurse, P. (1993). The fission yeast cdc18+ gene product couples S phase to START and mitosis. Cell 74, 371–382.

Kiang, L., Heichinger, C., Watt, S., Bahler, J., and Nurse, P. (2009). Cyclin-dependent kinase inhibits reinitiation of a normal S-phase program during G2 in fission yeast. Mol Cell Biol 29, 4025–4032.

Kitamura, K., Maekawa, H., and Shimoda, C. (1998). Fission yeast Ste9, a homolog of Hct1/Cdh1 and Fizzy-related, is a novel negative regulator of cell cycle progression during G1-phase. Mol Biol Cell 9, 1065–1080.

Lundgren, K., Walworth, N., Booher, R., Dembski, M., Kirschner, M., and Beach, D. (1991). mik1 and wee1 cooperate in the inhibitory tyrosine phosphorylation of cdc2. Cell 64, 1111–1122.

Millar, J.B., Lenaers, G., and Russell, P. (1992). Pyp3 PTPase acts as a mitotic inducer in fission yeast. EMBO J 11, 4933–4941.

Mondesert, O., McGowan, C.H., and Russell, P. (1996). Cig2, a B-type cyclin, promotes the onset of S in Schizosaccharomyces pombe. Mol Cell Biol 16, 1527–1533.

Moreno, S., Labib, K., Correa, J., and Nurse, P. (1994). Regulation of the cell cycle timing of Start in fission yeast by the rum1+ gene. J Cell Sci Suppl 18, 63–68.

Moreno, S., and Nurse, P. (1994). Regulation of progression through the G1 phase of the cell cycle by the rum1+ gene. Nature 367, 236–242.

Morgan, D.O. (2007). The Cell Cycle: Principles of Control. New Science Press: London.

Ng, S.S., Anderson, M., White, S., and McInerny, C.J. (2001). mik1(+) G1-S transcription regulates mitotic entry in fission yeast. FEBS Lett 503, 131–134.

Nishitani, H., Lygerou, Z., Nishimoto, T., and Nurse, P. (2000). The Cdt1 protein is required to license DNA for replication in fission yeast. Nature 404, 625–628.

Nishitani, H., and Nurse, P. (1995). p65cdc18 plays a major role controlling the initiation of DNA replication in fission yeast. Cell 83, 397–405.

Novak, B., and Tyson, J.J. (1997). Modeling the control of DNA replication in fission yeast. Proc Natl Acad Sci U S A 94, 9147–9152.

Novak, B., and Tyson, J.J. (2008). Design principles of biochemical oscillators. Nat Rev Mol Cell Biol 9, 981–991.

Nurse, P. (1975). Genetic control of cell size at cell division in yeast. Nature 256, 547–551.

Nurse, P. (1990). Universal control mechanism regulating onset of M-phase. Nature 344, 503–508.

Reymond, A., Marks, J., and Simanis, V. (1993). The activity of S.pombe DSC-1-like factor is cell cycle regulated and dependent on the activity of p34cdc2. EMBO J 12, 4325–4334.

Russell, P., and Nurse, P. (1986). cdc25+ functions as an inducer in the mitotic control of fission yeast. Cell 45, 145–153.

Russell, P., and Nurse, P. (1987). Negative regulation of mitosis by wee1+, a gene encoding a protein kinase homolog. Cell 49, 559–567.

Rustici, G., Mata, J., Kivinen, K., Lio, P., Penkett, C.J., Burns, G., Hayles, J., Brazma, A., Nurse, P., and Bahler, J. (2004). Periodic gene expression program of the fission yeast cell cycle. Nat Genet 36, 809–817.

Schmoller, K.M., Turner, J.J., Koivomagi, M., and Skotheim, J.M. (2015). Dilution of the cell cycle inhibitor Whi5 controls budding-yeast cell size. Nature 526, 268–272.

Stern, B., and Nurse, P. (1996). A quantitative model for the cdc2 control of S phase and mitosis in fission yeast. Trends Genet 12, 345–350.

Toda, T., Ochotorena, I., and Kominami, K. (1999). Two distinct ubiquitin-proteolysis pathways in the fission yeast cell cycle. Philos Trans R Soc Lond B Biol Sci 354, 1551–1557.

Tyson, J.J., Csikasz-Nagy, A., and Novak, B. (2002). The dynamics of cell cycle regulation. Bioessays 24, 1095–1109.

Tyson, J.J., and Novak, B. (2020). A Dynamical Paradigm for Molecular Cell Biology. Trends Cell Biol 30, 504–515.

Wood, E., and Nurse, P. (2015). Sizing up to divide: mitotic cell-size control in fission yeast. Annu Rev Cell Dev Biol 31, 11–29.

Wuarin, J., Buck, V., Nurse, P., and Millar, J.B. (2002). Stable association of mitotic cyclin B/Cdc2 to replication origins prevents endoreduplication. Cell 111, 419–431.

Yamaguchi, S., Murakami, H., and Okayama, H. (1997). A WD repeat protein controls the cell cycle and differentiation by negatively regulating Cdc2/B-type cyclin complexes. Mol Biol Cell 8, 2475–2486.

Yamaguchi, S., Okayama, H., and Nurse, P. (2000). Fission yeast Fizzy-related protein srw1p is a G(1)-specific promoter of mitotic cyclin B degradation. EMBO J 19, 3968–3977.

Zarzov, P., Decottignies, A., Baldacci, G., and Nurse, P. (2002). G(1)/S CDK is inhibited to restrain mitotic onset when DNA replication is blocked in fission yeast. EMBO J 21, 3370–3376.

