## Supplementary Text for "Computational modelling of chromosome re-replication in mutant strains of fission yeast"

##### *Mathematical model for the endoreplication of Cdc13-depleted strain*

Our core model describes the temporal dynamics of the four Cdc10-targeted gene-products by the following nonlinear ordinary differential equations (ODE):

$$\frac{d[\mathrm{Cig2}]_\mathrm{T}}{dt} = k_{\mathrm{s,cig2}} \cdot [\mathrm{Cdc10}] - k_{\mathrm{d,cig2}} \cdot [\mathrm{Cig2}]_\mathrm{T} \quad (1)$$

$$\frac{d[\mathrm{Nrm1}]_\mathrm{T}}{dt} = k_{\mathrm{s,nrm1}} \cdot [\mathrm{Cdc10}] - (k'_{\mathrm{d,nrm1}} + k_{\mathrm{d,nrm1}} \cdot [\mathrm{Ste9}]) \cdot [\mathrm{Nrm1}]_\mathrm{T} \quad (2)$$

$$\frac{d[\mathrm{Cdc18}]}{dt} = k_{\mathrm{s,cdc18}} \cdot [\mathrm{Cdc10}] - (k'_{\mathrm{d,cdc18}} + k_{\mathrm{d,cdc18}} \cdot [\mathrm{Cig2}]) \cdot [\mathrm{Cdc18}] \quad (3)$$

$$\frac{d[\mathrm{Mik1}]}{dt} = k_{\mathrm{s,mik1}} \cdot [\mathrm{Cdc10}] - \left(k'_{\mathrm{d,mik1}} + \frac{k_{\mathrm{d,mik1}}}{1+UDNA\cdot[\mathrm{Cdc18}]}\right) \cdot [\mathrm{Mik1}] \quad (4)$$

$\mathrm{Cig2_T}$ and $\mathrm{Nrm1_T}$ refer to total levels of Cdk1:Cig2 and Nrm1:Yox1 complexes, respectively, under the assumption that Cdk1 and Yox1 are plentiful. Nrm1, Cdc18 and Mik1 have both constitutive and regulated degradations. The degradation of Nrm1 and Cdc18 is activated by Ste9 and Cig2-kinase, respectively. In contrast, Mik1 degradation is inhibited by the unreplicated-DNA checkpoint (*UDNA*) in a Cdc18 concentration-dependent manner.

$[\mathrm{Cig2}]_\mathrm{T}$ represents the sum of dephosphorylated, active Cdk1:Cig2 complexes, [Cig2], and phosphorylated, inactive Cdk1P:Cig2 complexes, $[\mathrm{Cig2}]_\mathrm{T} - [\mathrm{Cig2}]$. The former is governed by the ODE:

$$\frac{d[\mathrm{Cig2}]}{dt} = k_{\mathrm{s,cig2}} \cdot [\mathrm{Cdc10}] - k_{\mathrm{p,cig2}} \cdot [\mathrm{Mik1}] \cdot [\mathrm{Cig2}] + k_{\mathrm{dp,cig2}} \cdot ([\mathrm{Cig2}]_\mathrm{T} - [\mathrm{Cig2}]) - k_{\mathrm{d,cig2}} \cdot [\mathrm{Cig2}] \quad (5)$$

where Cdk1:Cig2 is inactivated by phosphorylation by Mik1 and activated by an unspecified phosphatase whose constant concentration is lumped into the rate constant $k_{\mathrm{dp,cig2}}$.

Although the transcription of Ste9 is regulated by Cdc10 (Tournier and Millar, 2000), the fluctuation of its message is very small (Rustici *et al.*, 2004); so we assume constant total Ste9 concentration, $[\mathrm{Ste9}]_\mathrm{T}$. We use a Hill function to describe multi-site phosphorylation of Ste9 by Cdk1:Cig2 and Cdk1:Cdc13 complexes, because it happens on a fast time-scale. The expression for the unphosphorylated, active form of Ste9 is given by:

$$[\mathrm{Ste9}] = \frac{[\mathrm{Ste9}]_\mathrm{T} \cdot \alpha^n}{\alpha^n + ([\mathrm{Cig2}] + [\mathrm{Cdc13}])^n} \quad (6)$$

We assume that the cell volume-to-DNA ratio, $VpD$, increases exponentially with specific growth rate ($\mu$) for cell volume:

$$\frac{dVpD}{dt} = \mu \cdot VpD \quad (7)$$

and it is halved at DNA replication (i.e., when the rising Cig2-kinase activity crosses a threshold of 0.2).

In the model, we include a G1-specific inhibitor of Cdc10 transcription factor (TFI). In the 'titration model', we assume that TFI is present at constant concentration, $[\mathrm{TFI}]_\mathrm{T}$. The inactivation of TFI, which is described by a Hill function, is dependent on a starter kinase, SK, on Cdc13 activity, and on a product of Cdc10 transcription, which we equate to Cig2 protein level:

$$[\mathrm{TFI}]_{\mathrm{t}} = \frac{[\mathrm{TFI}]_{\mathrm{T}} \cdot \beta^m}{\beta^m \cdot (1 + VpD \cdot [\mathrm{SK}]) + ([\mathrm{Cig2}]_{\mathrm{T}} + [\mathrm{Cdc13}])^m} \tag{8}$$

where $\mathrm{TFI_t}$ represents free TFI and its complex with Cdc10, $[\mathrm{TFI}]_{\mathrm{t}}$ = [TFI] + [Cdc10:TFI].

In the 'TFI-dilution model', TFI is synthesized at a rate proportional to its gene-dosage:

$$\frac{\mathrm{d}[\mathrm{TFI}]_{\mathrm{T}}}{dt} = \frac{k_{\mathrm{s,tfi}}}{VpD} - k_{\mathrm{d,tfi}} \cdot [\mathrm{TFI}]_{\mathrm{T}} \quad . \tag{9}$$

In this case, $[\mathrm{TFI}]_{\mathrm{T}}$ is always moving toward a steady-state value, $k_{\mathrm{s,tfi}}/\left(k_{\mathrm{d,tfi}} \cdot VpD\right)$, which decreases exponentially as the cell grows and doubles abruptly in S phase when the *tfi* gene locus is replicated. There is no need for a starter kinase in the TFI-dilution model.

In our model, we assume that Nrm1 and TFI bind to Cdc10 independently resulting four different forms of Cdc10, and the total concentration of Cdc10, $[\mathrm{Cdc10}]_{\mathrm{T}}$ = [Cdc10] + [Nrm1:Cdc10] + [Cdc10:TFI] + [Nrm1:Cdc10:TFI], is assumed to be constant. The time rate of change for Nrm1-containing Cdc10 complexes, [Comp1] = [Nrm1:Cdc10] + [Nrm1:Cdc10:TFI], is given by:

$$\frac{d[\mathrm{Comp1}]}{dt} = k_{\mathrm{ass1}} \cdot ([\mathrm{Cdc10}]_{\mathrm{T}} - [\mathrm{Comp1}]) \cdot ([\mathrm{Nrm1}]_{\mathrm{T}} - [\mathrm{Comp1}]) - \left(k_{\mathrm{dis1}} + k'_{\mathrm{d,nrm1}} + k_{\mathrm{d,nrm1}} \cdot [\mathrm{Ste9}]\right) \cdot [\mathrm{Comp1}] \tag{10}$$

where [Nrm1] = $[\mathrm{Nrm1}]_{\mathrm{T}}$ – [Comp1]. If the association and dissociation rates are fast and $k_{\mathrm{dis1}} \gg k'_{\mathrm{d,nrm1}}$ and $k_{\mathrm{d,nrm1}}$[Ste9], then the Nrm1:Cdc10 complexes are in pseudo-steady state:

$$[\mathrm{Comp1}] = \frac{[\mathrm{Cdc10}]_{\mathrm{T}} + [\mathrm{Nrm1}]_{\mathrm{T}} + K_{\mathrm{dis1}} - \sqrt{([\mathrm{Cdc10}]_{\mathrm{T}} + [\mathrm{Nrm1}]_{\mathrm{T}} + K_{\mathrm{dis1}})^2 - 4 \cdot [\mathrm{Cdc10}]_{\mathrm{T}} \cdot [\mathrm{Nrm1}]_{\mathrm{T}}}}{2} \tag{11}$$

where $K_{\mathrm{dis1}} = k_{\mathrm{dis1}}/k_{\mathrm{ass1}}$. Similarly, the time rate of change of the TFI-containing Cdc10 complexes, [Comp5] = [Cdc10:TFI] + [Nrm1:Cdc10:TFI] = $[\mathrm{TFI}]_{\mathrm{T}}$ – [TFI], is given by:

$$\frac{d[\mathrm{Comp5}]}{dt} = k_{\mathrm{ass5}} \cdot ([\mathrm{Cdc10}]_{\mathrm{T}} - [\mathrm{Comp5}]) \cdot ([\mathrm{TFI}]_{\mathrm{T}} - [\mathrm{Comp5}]) - k_{\mathrm{dis5}} \cdot [\mathrm{Comp5}] \tag{12}$$

which in pseudo-steady state provides:

$$[\mathrm{Comp5}] = \frac{[\mathrm{Cdc10}]_{\mathrm{T}} + [\mathrm{TFI}]_{\mathrm{t}} + K_{\mathrm{dis5}} - \sqrt{([\mathrm{Cdc10}]_{\mathrm{T}} + [\mathrm{TFI}]_{\mathrm{t}} + K_{\mathrm{dis5}})^2 - 4 \cdot [\mathrm{Cdc10}]_{\mathrm{T}} \cdot [\mathrm{TFI}]_{\mathrm{t}}}}{2} \tag{13}$$

Let $K_{\mathrm{dis1}}$ (resp. $K_{\mathrm{dis5}}$) be the dissociation constant for Nrm1 (resp. TFI) binding to Cdc10. Then, since the binding steps are assumed to be independent:

$$[\mathrm{Nrm1:Cdc10}] \cdot [\mathrm{Cdc10:TFI}] = \frac{[\mathrm{Cdc10}]^2[\mathrm{Nrm1}][\mathrm{TFI}]}{K_{\mathrm{dN}} K_{\mathrm{dT}}} = [\mathrm{Cdc10}] \cdot [\mathrm{Nrm1{:}Cdc10{:}TFI}] \tag{14}$$

As a consequence, we can calculate free, active Cdc10, not bound to any of its inhibitors, by the algebraic equation:

$$[\mathrm{Cdc10}] = \frac{([\mathrm{Cdc10}]_{\mathrm{T}} - [\mathrm{Comp1}]) \cdot ([\mathrm{Cdc10}]_{\mathrm{T}} - [\mathrm{Comp5}])}{[\mathrm{Cdc10}]_{\mathrm{T}}} \tag{15}$$

Our dynamic simulations of this model can be easily reproduced by the freely available software XPP/AUTO (http://www.math.pitt.edu/~bard/xpp/xpp.html). The input to this tool is an '.ode' file, and

Fig. 4C can be reproduced using ‘Fig4C.ode’ provided in the Supplementary Files. To reproduce (C) panels on Figures 2-3 requires the parameter changes listed the following Table.

| **Parameters & their value on Fig.4C** | **Figure 2C** | **Figure 3C** |
|---|---|---|
| $k_{s,cig2}$ = 0.1 | 0.1 | 0.4 |
| $k_{d,cig2}$ = 0.05 | 0.05 | 0.2 |
| $\tau$ = 1 | 1 | 0.01 |
| $k_{s,mik1}$ = 1 | 1 | 0 |
| $k_{p,cig2}$ = 5 | 5 | 0 |
| $[TFI]_{tot}$ =1 | 0 | 1 |
| $n$ = 8 | 4 | 4 |

To simulate growth-controlled endoreplication by the titration or by the TFI-dilution model use ‘Figure 5C.ode’ or ‘Figure S1C.ode’, respectively.

***Phase plane analysis***

The dynamics of the network can be conveniently illustrated on an $Nrm1_T$ – $Cig2_T$ phase plane by using two ODE’s (Eqs. 1 and 2) and assuming pseudo-steady states for all other dynamic variables represented by Eqs. 3-5. By setting the right-hand side of these ODE’s to zero, the corresponding algebraic equations are used together with Eqs. 6, 8, 10, 12, 13 and 15 to fully describe the dynamical system.

Use ‘Figure 1C.ode’ to reproduce the phase plane on Fig. 1C. The ‘Core PP.ode’ file provides the phase plane diagram on Fig.4B and it can be used to calculate the phase planes of other figures after the following parameter changes:

| | **Required parameter changes in ‘Core PP.ode’ file** |
|---|---|
| **Fig.2B** | TFItot=0, n=4 |
| **Fig.3B** | kscig2=0.4, kdcig2=0.2, ksmik1=0, kpcig2=0, n=4 |
| **Fig.6B** | uDNA=1, Kdiss1=10 |
| **Fig.6D** | Cdc13=0.15 |
| **Fig.S1B** | TFItot=1.15, n=4 |

***Mathematical model for the protein overexpression strains***

In order to describe re-replication induced by overexpression of Rum1, Ste9 and Cdc18 in *cdc13*$^+$ genetic background, the core model has been modified and extended. An ODE for Cdk1:Cdc13 kinase has been introduced:

$$\frac{d[\mathrm{Cdc13}]}{dt} = k_{\mathrm{s,cdc13}} - (k'_{\mathrm{d,cdc13}} \cdot [\mathrm{Ste9}]_{\mathrm{T}} + k_{\mathrm{d,cdc13}} \cdot [\mathrm{Ste9}]) \cdot [\mathrm{Cdc13}] \qquad (16)$$

A constant rate of synthesis of Cdc13 is opposed by APC/C:Ste9-dependent degradation of Cdc13 cyclin. (We are assuming, as before, that Cdk1 is in excess over its cyclin partners.) The mitotic kinase is targeted to fast ($k_{d,cdc13}$) and slow ($k'_{d,cdc13}$) degradation by unphosphorylated and total Ste9, respectively.

The Cdc18 ODE (Eq.3) is supplemented by a constitutive synthesis term, $k'_{s,18}$, corresponding to synthesis from the Nmt-promoter:

$$\frac{d[\mathrm{Cdc18}]}{dt} = k'_{\mathrm{s},18} + k_{\mathrm{s},18} \cdot [\mathrm{Cdc10}] - (k'_{\mathrm{d},18} + k_{\mathrm{d},18} \cdot [\mathrm{Cig2}]) \cdot [\mathrm{Cdc18}] \qquad (17)$$

$k'_{s,18}$ takes values larger than zero in case of overexpression, i.e., in the absence of thiamine. Ste9 overexpression was modelled simply by increasing the parameter $[\mathrm{Ste9}]_T$ over its nominal value (0.5). To simplify the mathematical description, the Cdk1:cyclin inhibitor, Rum1, was also introduced as a parameter in the model. Besides Ste9-induced Cdc13 degradation, the three overexpressed proteins have an effect of the activities of Cdk1:Cyclin complexes. We assume that the inhibition of Cdk1:cyclin complexes by these inhibitory proteins is a fast and reversible process which can be described by equilibrium binding:

$$K_\mathrm{d} = \frac{[\mathrm{kinase}]_\mathrm{a} \cdot [\mathrm{Inhibitor}]}{[\mathrm{kinase}]_\mathrm{T} - [\mathrm{kinase}]_\mathrm{a}}$$

where $K_d$ is the equilibrium dissociation constant and the 'T' and 'a' subscripts refer to the total and the active form of the kinase. Using this concept, we calculate the active forms of Cdk1:Cdc13 and Cdk1:Cig2:

$$[\mathrm{Cdc13}]_\mathrm{a} = \frac{[\mathrm{Cdc13}]}{1+[\mathrm{Rum1}]+[\mathrm{Cdc18}]/K_{\mathrm{d18}}} \qquad (18)$$

$$[\mathrm{Cig2}]_\mathrm{a} = \frac{K_{\mathrm{d2}} \cdot [\mathrm{Cig2}]}{K_{\mathrm{d2}}+[\mathrm{Rum1}]+[\mathrm{Ste9}]_\mathrm{T}} \qquad (19)$$

Since Rum1 is a strong inhibitor of Cdk1:Cdc13 complex, we choose the corresponding $K_d$ value = 1, and $K_{d18}$ (=3) represents the strength of Cdc13-kinase inhibition by Cdc18 relative to Rum1. Rum1 is a weak inhibitor of Cdk1:Cig2, so we set $K_{d2}$ = 100. We also assume the Ste9 protein competitively inhibits Cig2-kinase with the same large dissociation constant. The catalytic action of Cig2- and Cdc13-kinase (Cig2 and Cdc13) in the core model are replaced by $\mathrm{Cig2}_a$ and $\mathrm{Cdc13}_a$ calculated by Eqs.18 & 19.

The phase planes, the bifurcation diagrams and time-course simulations for the overproducing strains can be reproduced by 'Figure 7.ode', 'Figure S2.ode' and 'Figure S3.ode', respectively.

**Supplementary references:**


Rustici, G., Mata, J., Kivinen, K., Lio, P., Penkett, C.J., Burns, G., Hayles, J., Brazma, A., Nurse, P., and Bahler, J. (2004). Periodic gene expression program of the fission yeast cell cycle. Nat Genet *36*, 809-817.

Tournier, S., and Millar, J.B. (2000). A role for the START gene-specific transcription factor complex in the inactivation of cyclin B and Cut2 destruction. Mol Biol Cell *11*, 3411-3424.

**Figure S1: Nucleocytoplasmic control of the oscillation.** (A) The effect of $TFI_T$ level (logarithmic scale) on the regulatory network. Small $TFI_T$ levels allow large amplitude oscillations (the filled circles are minima and maxima), but high levels of $TFI_T$ create stable steady states (solid line). Unstable steady states indicated by dashed lines. The black star shows the position of the system corresponding to panel B. (B) Higher than threshold level of $TFI_T$ creates a stable steady state near the origin. (C) Numerical simulation of the network shown on Fig.4A with nucleocytoplasmic control on TFI.

**Figure S2: Signal-Response (S-R) diagrams for overexpressing strains.** The levels of Cdc18 (A), Rum1 (B) and Ste9 (C) overexpression are expressed relative to endogenous levels in wild-type strain. Stable and unstable steady states are indicated by solid and dashed lines, respectively. The network oscillates around the unstable steady states with amplitudes between the filled circles.

**Figure S3: Temporal simulation of endoreplication cycles in Rum1 (A) and Ste9 (B) overexpression.** Observe that the cytoplasmic mass per DNA ratio is higher in $rum1^{op}$ than in $ste9^{op}$ cells.

Figure S1

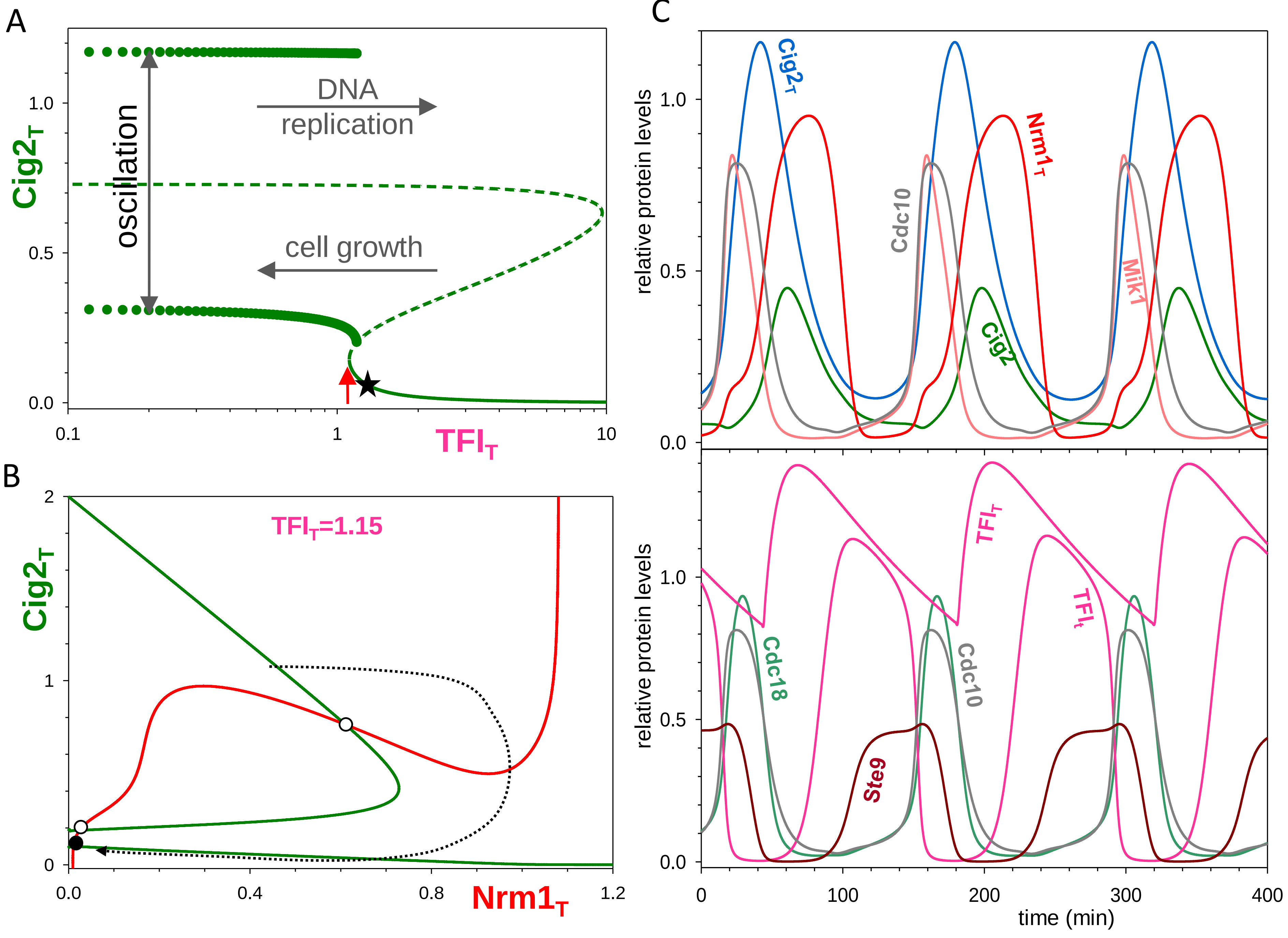
A
Cig2T
1.0
0.5
0.0
0.1
1
10
TFIT
oscillation
DNA
replication
cell growth
B
Cig2T
2
1
0
0.0
0.4
0.8
1.2
Nrm1T
TFIT=1.15
C
relative protein levels
Cig2T
Nrm1T
Cdc10
Mik1
Cig2
1.0
0.5
0.0
relative protein levels
TFIT
TFIt
Cdc18
Cdc10
Ste9
1.0
0.5
0.0
0
100
200
300
400
time (min)

Figure S2

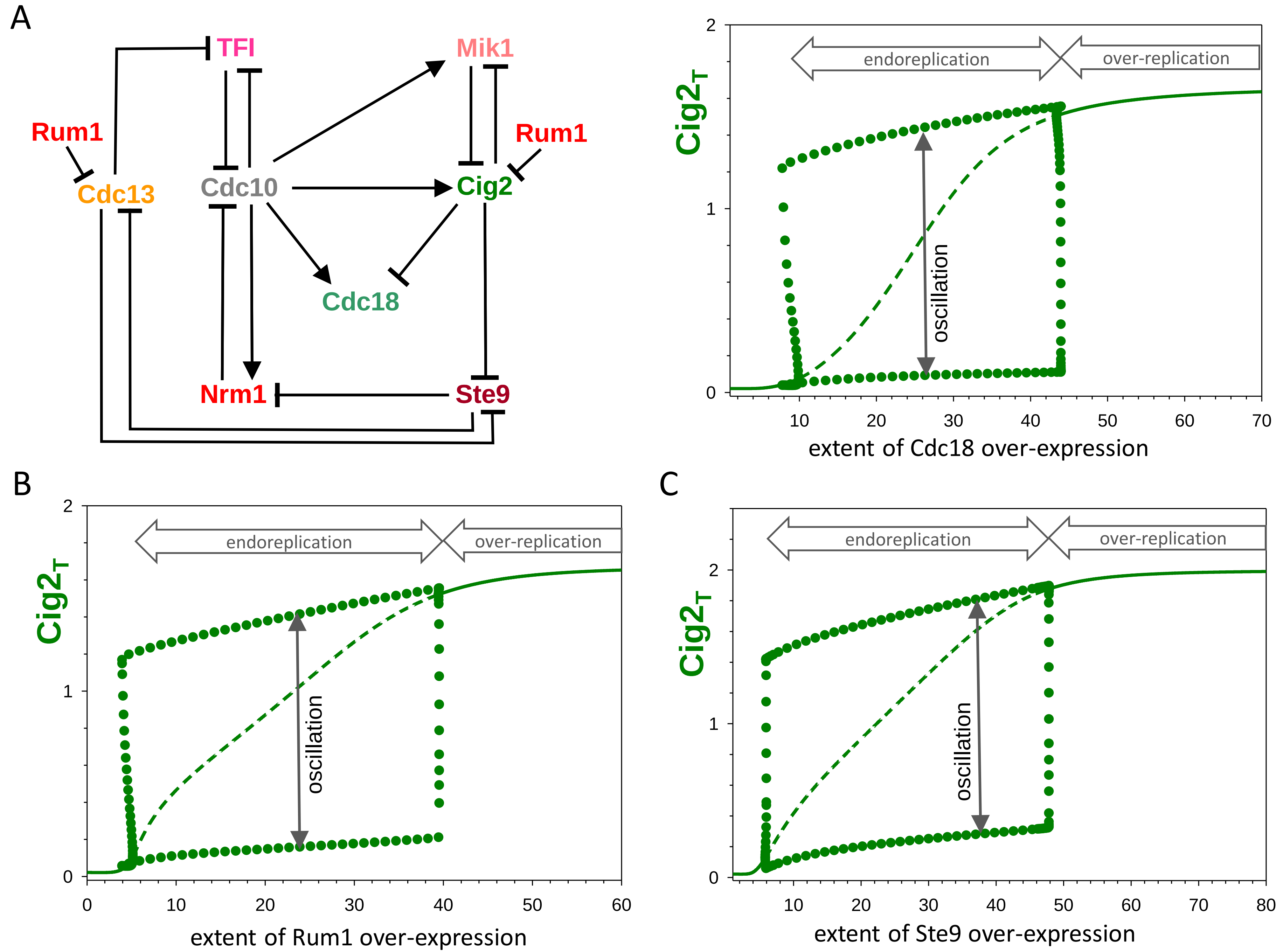
A
TFI
Mik1
Rum1
Rum1
Cdc13
Cdc10
Cig2
Cdc18
Nrm1
Ste9
endoreplication
over-replication
oscillation
Cig2T
extent of Cdc18 over-expression
B
endoreplication
over-replication
oscillation
Cig2T
extent of Rum1 over-expression
C
endoreplication
over-replication
oscillation
Cig2T
extent of Ste9 over-expression

Figure S3

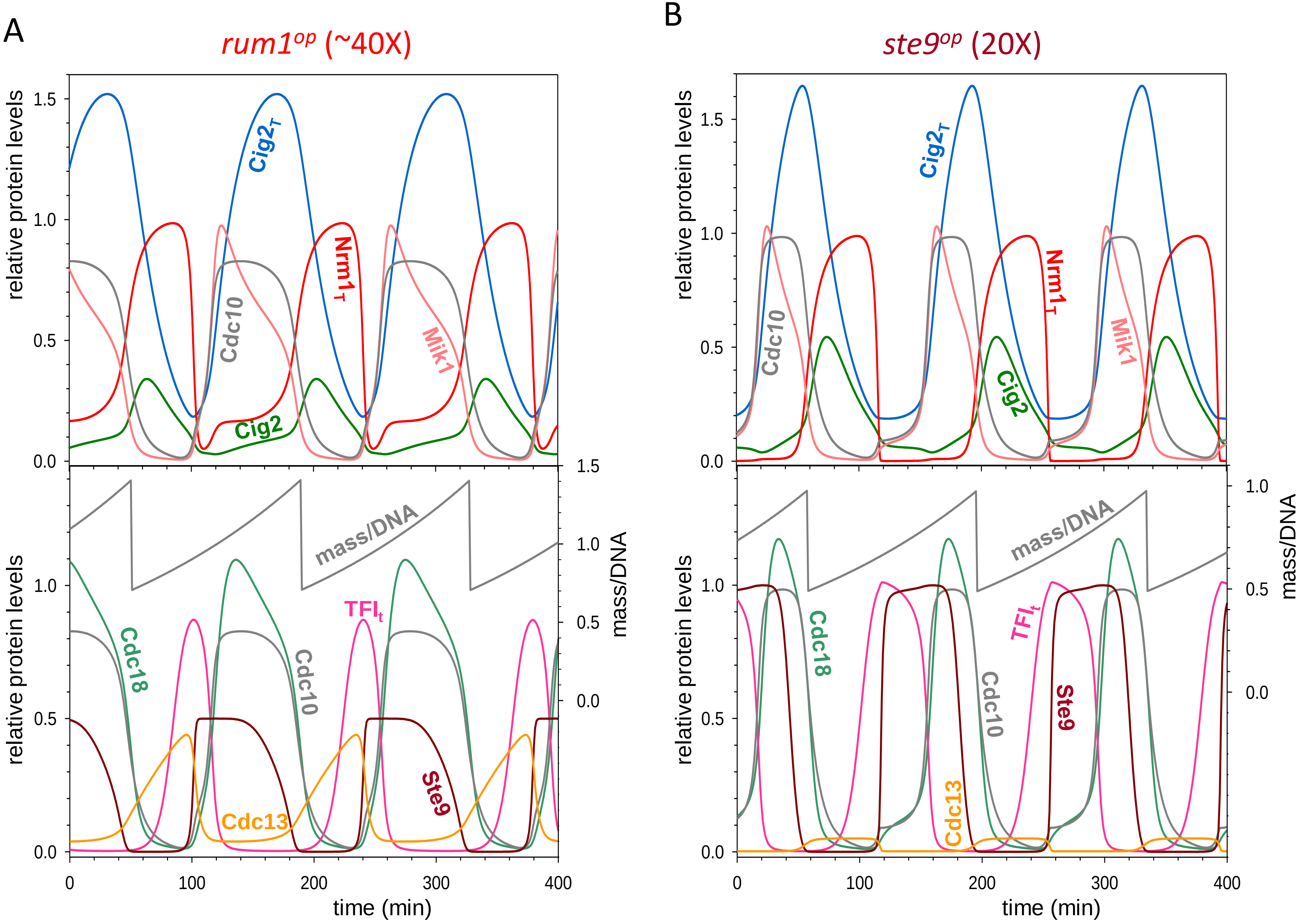
A
rum1op (~40X)
B
ste9op (20X)
relative protein levels
mass/DNA
time (min)
Cig2T
Nrm1T
Cdc10
Mik1
Cig2
Cdc18
TFIt
Ste9
Cdc13

### XPP/AUTO Models

Model simulations were performed using the freely available software XPP/AUTO (http://www.math.pitt.edu/~bard/xpp/xpp.html). The models are provided below in the form of .ode files readable by XPP. To use, copy the code into a text editor and save as [filename].ode.

**Figure 1C**

```
# .ode file for Figure 1C
# Differential equations
Cig2'  = kscig2*Cdc10 - kdcig2*Cig2
Nrm1T' = ksnrm1*Cdc10 - (kdnrm1' + kdnrm1*Ste9)*Nrm1T
# Algebraic equations
Ste9 = Ste9T*alpha^n/(alpha^n + Cig2^n)
BB1 = Cdc10T + Nrm1T + Kdiss1
Comp1 = (BB1 - sqrt(BB1^2 - 4*Cdc10T*Nrm1T))/2
Cdc10 = Cdc10T-Comp1
# Parameter values
p kscig2=0.1, kdcig2=0.05, ksnrm1=1, kdnrm1'=0.01, kdnrm1=10
p Ste9T=0.5, alpha=0.1, n=4, Cdc10T=1, Kdiss1=0.001
# XPP settings
@ METH=stiff, XLO=0,XHI=1.2,YLO=0,YHI=2,NMESH=400, XP=Nrm1T, YP=Cig2
done
```

**Core PP**

```
# XPP/AUTO .ode file for phaseplane calculations
# Differential equations
Cig2T' = kscig2*Cdc10 - kdcig2*Cig2T
Nrm1T' = ksnrm1*Cdc10 - (kdnrm1' + kdnrm1*Ste9)*Nrm1T
# Algebraic equations
Cig2 = (kscig2*Cdc10 + kdpcig2*Cig2T)/(kpcig2*Mik1 + kdpcig2 + kdcig2)
Cdc18  = kscdc18*Cdc10/(kdcdc18' + kdcdc18"*Cig2)
Mik1 = ksmik1*Cdc10/(kdmik1' + kdmik1"*Cig2/(1 + uDNA*Cdc18))
Ste9 = Ste9T*alpha^n/(alpha^n + (Cig2 + Cdc13)^n)
BB1 = Cdc10T + Nrm1T + Kdiss1
Comp1 = (BB1 - sqrt(BB1^2 - 4*Cdc10T*Nrm1T))/2
TFIt = TFItot*beta^m/(beta^m*(1+SK) + (Cig2T + Cdc13)^m)
BB5= TFIt + Cdc10T + Kdiss5
Comp5 = (BB5 - sqrt(BB5^2 - 4*TFIt*Cdc10T))/2
Cdc10 = (Cdc10T-Comp1)*(Cdc10T-Comp5)/Cdc10T
# Parameter values
p kscig2=0.1, kdcig2=0.05, kpcig2=5, kdpcig2=0.2, kscdc18=0.2, kdcdc18'=0.1, kdcdc18"=1
p ksnrm1=1, kdnrm1'=0.01, kdnrm1=10, tau=1, ksmik1=1, kdmik1'=0.5, kdmik1"=10, uDNA=0
p Ste9T=0.5, alpha=0.1, n=8, Cdc10T=1, Kdiss1=0.001
p TFItot=1, beta=0.3, m=4, Kdiss5=0.01, Cdc13=0, SK=0
# XPP settings
@ METH=stiff, XLO=0, XHI=1.2, YLO=0, YHI=2, NMESH=400, total=200, dt=1
@ XP=Nrm1T, YP=Cig2T
done
```

**Figure 4C**

```
# .ode file for simulation of Figure 2C,3C & 4C
# Differential equations
Cig2T'   = kscig2*Cdc10 - kdcig2*Cig2T
Cig2'    = kscig2*Cdc10 - kpcig2*Mik1*Cig2 + kdpcig2*(Cig2T-Cig2) - kdcig2*Cig2
Cdc18'  = kscdc18*Cdc10 - (kdcdc18'+kdcdc18"*Cig2)*Cdc18
Nrm1T' = (ksnrm1*Cdc10 - (kdnrm1' + kdnrm1*Ste9)*Nrm1T)*tau
Mik1' = ksmik1*Cdc10 - (kdmik1' + kdmik1"*Cig2/(1 + uDNA*Cdc18))*Mik1
# Algebraic equations
Ste9 = Ste9T*alpha^n/(alpha^n + (Cig2 + Cdc13)^n)
BB1 = Cdc10T + Nrm1T + Kdiss1
Comp1 = (BB1 - sqrt(BB1^2 - 4*Cdc10T*Nrm1T))/2
TFIt = TFItot*beta^m/(beta^m + (Cig2T + Cdc13)^m)
BB5= TFIt + Cdc10T + Kdiss5
Comp5 = (BB5 - sqrt(BB5^2 - 4*TFIt*Cdc10T))/2
Cdc10 = (Cdc10T-Comp1)*(Cdc10T-Comp5)/Cdc10T
# Auxiliary variables
aux TFIt = TFItot*beta^m/(beta^m + (Cig2T + Cdc13)^m)
aux Cdc10 = (Cdc10T-Comp1)*(Cdc10T-Comp5)/Cdc10T
aux Ste9 = Ste9T*alpha^n/(alpha^n + (Cig2 + Cdc13)^n)
# Parameter values
p kscig2=0.1, kdcig2=0.05, kpcig2=5, kdpcig2=0.2
p kscdc18=0.2, kdcdc18'=0.1, kdcdc18"=1
p ksnrm1=1, kdnrm1'=0.01, kdnrm1=10, tau=1
p ksmik1=1, kdmik1'=0.5, kdmik1"=10, uDNA=0
p Ste9T=0.5, alpha=0.1, n=8
p Cdc10T=1, Kdiss1=0.001
p TFItot=1, beta=0.3, m=4, Kdiss5=0.01, Cdc13=0
# XPP settings
@ METH=stiff, XLO=0, XHI=150, YLO=0, YHI=1.2, total=180, dt=1
@ XP=time, nplot=8, YP=Cig2T, yp2=Cig2, yp3=Cdc18, yp4=Mik1
@ yp5=Nrm1T, yp6=Ste9, yp7=TFIt, yp8=Cdc10
done
```

**Figure 5C**

```
# .ode file for simulation of Figure 5C
# Differential equations
Cig2T'   = kscig2*Cdc10 - kdcig2*Cig2T
Cig2'    = kscig2*Cdc10 - kpcig2*Mik1*Cig2 + kdpcig2*(Cig2T-Cig2) - kdcig2*Cig2
Cdc18'  = kscdc18*Cdc10 - (kdcdc18'+kdcdc18"*Cig2)*Cdc18
Nrm1T' = (ksnrm1*Cdc10 - (kdnrm1' + kdnrm1*Ste9)*Nrm1T)*tau
Mik1' = ksmik1*Cdc10 - (kdmik1' + kdmik1"*Cig2/(1 + uDNA*Cdc18))*Mik1
VpD' = mu*VpD
# Algebraic equations
Ste9 = Ste9T*alpha^n/(alpha^n + (Cig2 + Cdc13)^n)
BB1 = Cdc10T + Nrm1T + Kdiss1
Comp1 = (BB1 - sqrt(BB1^2 - 4*Cdc10T*Nrm1T))/2
TFIt = TFItot*beta^m/(beta^m*(1+SK*VpD) + (Cig2T + Cdc13)^m)
BB5= TFIt + Cdc10T + Kdiss5
Comp5 = (BB5 - sqrt(BB5^2 - 4*TFIt*Cdc10T))/2
Cdc10 = (Cdc10T-Comp1)*(Cdc10T-Comp5)/Cdc10T
# Auxiliary variables
aux TFIt = TFItot*beta^m/(beta^m*(1+SK*VpD) + (Cig2T + Cdc13)^m)
aux Cdc10 = (Cdc10T-Comp1)*(Cdc10T-Comp5)/Cdc10T
aux Ste9 = Ste9T*alpha^n/(alpha^n + (Cig2 + Cdc13)^n)
global +1 {cig2-thres} {VpD=VpD/2}
# Initial values
init Cig2T=0.94,Cig2=0.40,Cdc18=0.11,Nrm1T=0.79,Mik1=0.04,VpD=0.52
# Parameter values
p kscig2=0.1, kdcig2=0.05, kpcig2=5, kdpcig2=0.2
p kscdc18=0.2, kdcdc18'=0.1, kdcdc18"=1
p ksnrm1=1, kdnrm1'=0.01, kdnrm1=10, tau=0.1
p ksmik1=1, kdmik1'=0.5, kdmik1"=10, uDNA=0
p Ste9T=0.5, alpha=0.1, n=8
p Cdc10T=1, Kdiss1=0.001
p TFItot=2,beta=0.3, m=4, Kdiss5=0.01
p Cdc13=0, SK=1.25, mu=0.005, thres=0.2
# XPP settings
@ METH=stiff,XLO=0,XHI=400,YLO=0,YHI=1.2,total=400,dt=1,bounds=1000
@ XP=time,nplot=8,YP=Cig2T,yp2=Cig2,yp3=Cdc18,yp4=Nrm1T
@ yp5=Mik1,yp6=TFItot,yp7=TFIt,yp8=Cdc10
done
```

**Figure S1C**

```
# .ode file for simulation of Figure S1
# Differential equations
Cig2T'   = kscig2*Cdc10 - kdcig2*Cig2T
Cig2'    = kscig2*Cdc10 - kpcig2*Mik1*Cig2 + kdpcig2*(Cig2T-Cig2) - kdcig2*Cig2
Cdc18'  = kscdc18*Cdc10 - (kdcdc18'+kdcdc18"*Cig2)*Cdc18
Nrm1T' = (ksnrm1*Cdc10 - (kdnrm1' + kdnrm1*Ste9)*Nrm1T)*tau
Mik1' = ksmik1*Cdc10 - (kdmik1' + kdmik1"*Cig2/(1 + uDNA*Cdc18))*Mik1
TFItot' = kstfi/VpD - kdtfi*TFItot
VpD' = mu*VpD
# Algebraic equations
Ste9 = Ste9T*alpha^n/(alpha^n + (Cig2 + Cdc13)^n)
BB1 = Cdc10T + Nrm1T + Kdiss1
Comp1 = (BB1 - sqrt(BB1^2 - 4*Cdc10T*Nrm1T))/2
TFIt = TFItot*beta^m/(beta^m + (Cig2T + Cdc13)^m)
BB5= TFIt + Cdc10T + Kdiss5
Comp5 = (BB5 - sqrt(BB5^2 - 4*TFIt*Cdc10T))/2
Cdc10 = (Cdc10T-Comp1)*(Cdc10T-Comp5)/Cdc10T
# Auxiliary variables
aux TFIt = TFItot*beta^m/(beta^m + (Cig2T + Cdc13)^m)
aux Cdc10 = (Cdc10T-Comp1)*(Cdc10T-Comp5)/Cdc10T
aux Ste9 = Ste9T*alpha^n/(alpha^n + (Cig2 + Cdc13)^n)
global +1 {cig2-thres} {VpD=VpD/2}
# Initial values
init VpD=1,Cig2T=0.Cig2=1,Cdc18=0,Nrm1T=0,Mik1=0,TFItot=1
# Parameter values
p kscig2=0.1, kdcig2=0.05, kpcig2=5, kdpcig2=0.2
p kscdc18=0.2, kdcdc18'=0.1, kdcdc18"=1
p ksnrm1=1, kdnrm1'=0.01, kdnrm1=10, tau=0.1
p ksmik1=1, kdmik1'=0.5, kdmik1"=10, uDNA=0
p Ste9T=0.5, alpha=0.1, n=4
p Cdc10T=1, Kdiss1=0.001
p beta=0.3, m=4, Kdiss5=0.01, Cdc13=0
p kstfi=0.1, kdtfi=0.1, mu=0.005, thres=0.2
# XPP settings
@ METH=stiff,XLO=0,XHI=1000,YLO=0,YHI=1.4,total=1000,dt=1,bounds=1000
@ XP=time,nplot=8,YP=Cig2T,yp2=Cig2,yp3=Cdc18,yp4=Nrm1T
@ yp5=Mik1,yp6=TFItot,yp7=TFIt,yp8=Cdc10
done
```

**Figure 7**

```
# XPP/AUTO .ode file for phaseplane calculations on Figure 7
# Differential equations
Cig2T' = kscig2*Cdc10 - kdcig2*Cig2T
Nrm1T' = ksnrm1*Cdc10 - (kdnrm1' + kdnrm1*Ste9)*Nrm1T
# Algebraic equations
Cig2 = (kscig2*Cdc10 + kdpcig2*Cig2T)/(kpcig2*Mik1 + kdpcig2 + kdcig2)
Cig2a = Cig2*Kdiss2/(Kdiss2 + Rum1 + Ste9T)
Cdc18  = (kscdc18' + kscdc18*Cdc10)/(kdcdc18' + kdcdc18"*Cig2a)
Mik1 = ksmik1*Cdc10/(kdmik1' + kdmik1"*Cig2a/(1 + uDNA*Cdc18))
Cdc13 = kscdc13/(kdcdc13'*Ste9T + kdcdc13*Ste9)
Cdc13a = Cdc13/(1+Rum1+Cdc18/Kd18)
Ste9 = Ste9T*alpha^n/(alpha^n + (Cig2a + Cdc13a)^n)
BB1 = Cdc10T + Nrm1T + Kdiss1
Comp1 = (BB1 - sqrt(BB1^2 - 4*Cdc10T*Nrm1T))/2
TFIt = TFItot*beta^m/(beta^m*(1+SK) + (Cig2T + Cdc13a)^m)
BB5= TFIt + Cdc10T + Kdiss5
Comp5 = (BB5 - sqrt(BB5^2 - 4*TFIt*Cdc10T))/2
Cdc10 = (Cdc10T-Comp1)*(Cdc10T-Comp5)/Cdc10T
# Parameter values
p kscig2=0.1, kdcig2=0.05, kpcig2=5, kdpcig2=0.2
p kscdc18'=0, kscdc18=0.2, kdcdc18'=0.1, kdcdc18"=1
p ksnrm1=1, kdnrm1'=0.01, kdnrm1=10, tau=1
p ksmik1=1, kdmik1'=0.5, kdmik1"=10, uDNA=0
p Ste9T=0.5, alpha=0.1, n=8
p Cdc10T=1, Kdiss1=0.001
p TFItot=1.1, beta=0.3, m=4, Kdiss5=0.01, SK=0, Rum1=0, Kdiss2=100, Kd18=3
p kscdc13=0.01, kdcdc13'=0.02, kdcdc13=0.5
# XPP settings
@ METH=stiff, XLO=0, XHI=1.2, YLO=0, YHI=2, NMESH=400, total=200, dt=1
@ XP=Nrm1T, YP=Cig2T
done
```

Comments:
panel B: 50-fold Cdc18 overexpression use kscdc18'=10 (=50*kscdc18) and uDNA=0.02,
panel C: 30-fold Rum1 overexpression use Rum1=30,
panel D: 20-fold Ste9 overexpression use Ste9t=10 (=20*0,5).

**Figure S2**

```
# .ode file for simulation of bifurcation diagrams on Figure S2
# Differential equations
Cig2T'   = kscig2*Cdc10 - kdcig2*Cig2T
Cig2'    = kscig2*Cdc10 - kpcig2*Mik1*Cig2 + kdpcig2*(Cig2T-Cig2) - kdcig2*Cig2
Cig2a' = Kdiss2*(Cig2-Cig2a) - Cig2a*(Rum1+Ste9T)
Cdc18'  = kscdc18' + kscdc18*Cdc10 - (kdcdc18'+kdcdc18"*Cig2a)*Cdc18
Nrm1T' = (ksnrm1*Cdc10 - (kdnrm1' + kdnrm1*Ste9)*Nrm1T)*tau
Mik1' = ksmik1*Cdc10 - (kdmik1' + kdmik1"*Cig2a/(1 + uDNA*Cdc18))*Mik1
Cdc13' = kscdc13 - (kdcdc13'*Ste9T + kdcdc13*Ste9)*Cdc13
# Algebraic equations
Cdc13a = Cdc13/(1+Rum1+Cdc18/Kd18)
Ste9 = Ste9T*alpha^n/(alpha^n + (Cig2a + Cdc13a)^n)
BB1 = Cdc10T + Nrm1T + Kdiss1
Comp1 = (BB1 - sqrt(BB1^2 - 4*Cdc10T*Nrm1T))/2
TFIt = TFItot*beta^m/(beta^m*(1+SK*VpD) + (Cig2T + Cdc13a)^m)
BB5= TFIt + Cdc10T + Kdiss5
Comp5 = (BB5 - sqrt(BB5^2 - 4*TFIt*Cdc10T))/2
Cdc10 = (Cdc10T-Comp1)*(Cdc10T-Comp5)/Cdc10T
# Auxiliary variables
aux TFIt = TFItot*beta^m/(beta^m*(1+SK*VpD) + (Cig2T + Cdc13a)^m)
aux Cdc10 = (Cdc10T-Comp1)*(Cdc10T-Comp5)/Cdc10T
aux Ste9 = Ste9T*alpha^n/(alpha^n + (Cig2a + Cdc13a)^n)
# Initial values
initial Cig2T=0, Cig2=0, Cig2a=0, Cdc18=0, Nrm1T=1, Mik1=0, Cdc13=1
# Parameter values
p Rum1=0, Ste9T=0.5, kscdc18'=0
p kscig2=0.1, kdcig2=0.05, kpcig2=5, kdpcig2=0.2
p kscdc18=0.2, kdcdc18'=0.1, kdcdc18"=1
p ksnrm1=1, kdnrm1'=0.01, kdnrm1=10, tau=0.1
p ksmik1=1, kdmik1'=0.5, kdmik1"=10, uDNA=0.02
p alpha=0.1, n=8
p Cdc10T=1, Kdiss1=0.001
p TFItot=1, beta=0.3, m=4, Kdiss5=0.01
p SK=0, VpD=0, mu=0.005, thres=0.2, Kdiss2=100, Kd18=3
p kscdc13=0.01, kdcdc13'=0.02, kdcdc13=0.5
# XPP settings
@ Method=stiff, Total=400, Bounds=100, Dt=1
@ Xplot=t, YPlot=Cig2T, Xlo=0, Xhi=400, Ylo=0, Yhi=2
@ NTST=150,NMAX=1000000,NPR=10000,DS=0.01,BOUNDS=2000
@ DSMAX=0.1,DSMIN=0.01,PARMIN=0,PARMAX=70, AUTOVAR=Cig2T
@ AUTOXMIN=0,AUTOXMAX=70,AUTOYMIN=0,AUTOYMAX=2
done
```

Comments: The code in its present form calculates the bifurcation diagram for Rum1 overexpression on Fig.S2B. In order to calculate the diagram for Cdc18 and Ste9 overexpression, Ste9t and kscdc18’ should be at first place in the parameter list. The parameter range (PARMAX and AUTOXMAX) has to be changed to 14 and 40 in case of panel A and panel D, respectively. For *cdc18*$^{op}$ also set uDNA to 0.02.

**Figure S3**

```
# .ode file for simulation of Figure S3
# Differential equations
Cig2T'   = kscig2*Cdc10 - kdcig2*Cig2T
Cig2'    = kscig2*Cdc10 - kpcig2*Mik1*Cig2 + kdpcig2*(Cig2T-Cig2) - kdcig2*Cig2
Cig2a' = Kdiss2*(Cig2-Cig2a) - Cig2a*(Rum1+Ste9T)
Cdc18'  = kscdc18' + kscdc18*Cdc10 - (kdcdc18'+kdcdc18"*Cig2a)*Cdc18
Nrm1T' = (ksnrm1*Cdc10 - (kdnrm1' + kdnrm1*Ste9)*Nrm1T)*tau
Mik1' = ksmik1*Cdc10 - (kdmik1' + kdmik1"*Cig2a/(1 + uDNA*Cdc18))*Mik1
Cdc13' = kscdc13 - (kdcdc13'*Ste9T + kdcdc13*Ste9)*Cdc13
VpD' = mu*VpD
# Algebraic equations
Cdc13a = Cdc13/(1+Rum1+Cdc18/Kd18)
Ste9 = Ste9T*alpha^n/(alpha^n + (Cig2a + Cdc13a)^n)
BB1 = Cdc10T + Nrm1T + Kdiss1
Comp1 = (BB1 - sqrt(BB1^2 - 4*Cdc10T*Nrm1T))/2
TFIt = TFItot*beta^m/(beta^m*(1+SK*VpD) + (Cig2T + Cdc13a)^m)
BB5= TFIt + Cdc10T + Kdiss5
Comp5 = (BB5 - sqrt(BB5^2 - 4*TFIt*Cdc10T))/2
Cdc10 = (Cdc10T-Comp1)*(Cdc10T-Comp5)/Cdc10T
# Auxiliary variables
aux TFIt = TFItot*beta^m/(beta^m*(1+SK*VpD) + (Cig2T + Cdc13a)^m)
aux Cdc10 = (Cdc10T-Comp1)*(Cdc10T-Comp5)/Cdc10T
aux Ste9 = Ste9T*alpha^n/(alpha^n + (Cig2a + Cdc13a)^n)
aux Cdc13a = Cdc13/(1+Rum1+Cdc18/Kd18)
global +1 {cig2a-thres} {VpD=VpD/2}
# Initial values
init Cig2T=0.94,Cig2=0.40,Cdc18=0.11,Nrm1T=0.79,Mik1=0.04,VpD=0.52
# Parameter values
p kscig2=0.1, kdcig2=0.05, kpcig2=5, kdpcig2=0.2
p kscdc18'=0, kscdc18=0.2, kdcdc18'=0.1, kdcdc18"=1
p ksnrm1=1, kdnrm1'=0.01, kdnrm1=10, tau=0.1
p ksmik1=1, kdmik1'=0.5, kdmik1"=10, uDNA=0
p Ste9T=0.5, alpha=0.1, n=8
p Cdc10T=1, Kdiss1=0.001
p TFItot=2,beta=0.3, m=4, Kdiss5=0.01
p SK=1.25, mu=0.005, thres=0.2, Rum1=0, Kdiss2=100, Kd18=3
p kscdc13=0.01, kdcdc13'=0.02, kdcdc13=0.5
# XPP settings
@ METH=stiff,XLO=0,XHI=400,YLO=0,YHI=1.6,total=400,dt=1,bounds=1000
@ XP=time,nplot=8,YP=Cig2T,yp2=Cig2a,yplot3=Cdc18,yp4=Nrm1T
@ yp5=Vpd,yp6=TFItot,yp7=TFIt,yp8=Cdc10
done
```

Comments: use Rum1=37 for panel A and Ste9t=10 (=20*0.5).
